# Multifunctionality of belowground food webs: resource, size and spatial energy channels

**DOI:** 10.1101/2021.06.06.447267

**Authors:** Anton Potapov

## Abstract

The belowground compartment of terrestrial ecosystems drives nutrient cycling, the decomposition and stabilisation of organic matter, and supports aboveground life. Belowground consumers create complex food webs that regulate functioning, ensure stability and support biodiversity both below and above ground. However, existing soil food-web reconstructions do not match recently accumulated empirical evidence and there is no comprehensive reproducible approach that accounts for the complex resource, size and spatial structure of food webs in soil. Here I build on generic food-web organization principles and use multifunctional classification of soil protists, invertebrates and vertebrates, to reconstruct ‘multichannel’ food-web across size classes of soil-associated consumers. This reconstruction is based on overlying feeding preference, prey protection, size spectrum and spatial distribution matrices combined with biomasses of trophic guilds to infer weighted trophic interactions. I then use food-web reconstruction, together with assimilation efficiencies, to calculate energy fluxes assuming a steady-state energetic system. Based on energy fluxes, I describe a number of indicators, related to stability, biodiversity and multiple ecosystem-level functions such as herbivory, top-down control, translocation and transformation of organic matter. I illustrate the approach with an empirical example, comparing it with traditional resource-focused soil food-web reconstruction. The multichannel reconstruction can be used to assess trophic multifunctionality (analogous to ecosystem multifunctionality), i.e. simultaneous support of multiple trophic functions by the food-web, and compare it across communities and ecosystems spanning beyond the soil. With further validation and parametrization, my multichannel reconstruction approach provides an effective tool for understanding and analysing soil food webs. I believe that having this tool will inspire more people to comprehensively describe soil communities and belowground-aboveground interactions. Such studies will provide informative indicators for including consumers as active agents in biogeochemical models, not only locally but also on regional and global scales.

## I. Introduction

### (1) Belowground communities and ecosystem functioning

Belowground communities regulate the decomposition and sequestration of organic matter in terrestrial ecosystems. Because they are responsible for processing a major part of primary production (Cebrian, 1999), the detrital system plays a central role in the carbon cycle (Clemmensen *et al*., 2013; Averill, Turner, & Finzi, 2014; Crowther *et al*., 2019), nitrogen cycle (Li *et al*., 2019) and other biogeochemical cycles (Xu, Thornton, & Post, 2013; Crowther *et al*., 2019). Microorganisms carry out the basic soil ecosystem processes.

Nevertheless, consumers of microorganisms and plant materials, have strong indirect effects on these processes by microbial grazing, litter shredding, organic matter transformation and translocation (Lavelle *et al*., 1997; Briones, 2014; Filser *et al*., 2016; Thakur & Geisen, 2019). However, the direction and magnitude of consumer effects are context-dependent and hard to predict and thus common global biogeochemical models often simply ignore them (Deckmyn *et al*., 2020). In recent years, considerable progress has been made in understanding the global distribution patterns of soil consumers (Crowther *et al*., 2019; Phillips *et al*., 2019; van den Hoogen *et al*., 2019; Potapov *et al*., 2020; Oliverio *et al*., 2020; Guerra *et al*., 2020) and calls to account for soil fauna in global biogeochemical models are becoming increasingly common (Filser *et al*., 2016; Grandy *et al*., 2016; Soong & Nielsen, 2016; Deckmyn *et al*., 2020). Some models have already been suggested (Chertov *et al*., 2017; Deckmyn *et al*., 2020). However, the usability of any models developed depends largely on whether or not they correctly depict the key functional groups and their interactions in soil.

### (2) Holistic approach to describe consumer community in soil

Soil communities include a huge diversity of consumers that spans phyla, size classes, trophic levels, and vertical layers (Anderson, 1975; Swift, Heal, & Anderson, 1979; Coleman, Callaham, & Crossley Jr, 2017; Potapov *et al*., 2021b). And soil ecosystem functioning is jointly driven by multiple components of the soil biota, including microorganisms, micro-meso- and macrofauna (Bradford *et al*., 2002; Wagg *et al*., 2014; Delgado-Baquerizo *et al*., 2020). This functional complexity calls for a holistic approach to describing soil communities across consumers of different body sizes, similar to the size spectrum approach commonly used in marine ecosystems (Blanchard *et al*., 2017). There were several conceptual and empirical attempts to apply the size spectrum approach to terrestrial belowground communities (Mulder, 2006; Petchey & Belgrano, 2010; Turnbull, George, & Lindo, 2014), but they provide only simplified information because terrestrial food webs have more complex size structures than marine ones (Potapov *et al*., 2019a, 2021b). The food-web framework is, however, a promising way of describing the functioning of terrestrial food webs because it unites the functional, biodiversity and stability aspects of biological systems (Hines *et al*., 2015; Barnes *et al*., 2018). Indeed, soil food-web properties may explain various soil functions better than environmental variation alone (de Vries *et al*., 2013).

### (3) Belowground food-web reconstructions

Most studies exploring the functioning of soil food webs assume the dominant role of basal resources in structuring food-web topology, stemming from the seminal work of William Hunt and colleagues (Hunt *et al*., 1987). These ‘traditional’ resource-based reconstructions were used to estimate energy fluxes and quantify nitrogen mineralisation in bacterial, fungal and plant energy channels in grasslands and agroecosystems (Hunt *et al*., 1987; de Ruiter *et al*., 1993). The approach has been also used to explore patterns of interaction strengths and was developed into the concept of ‘fast’ (e.g. bacterial) versus ‘slow’ (e.g. fungal) energy channels, jointly driving ecosystem stability (de Ruiter, Neutel, & Moore, 1995; Rooney *et al*., 2006). However, these ideas were mostly conceptualised for, and applied in, micro-food webs (protists, nematodes, microarthropods) in soil (Moore, McCann, & de Ruiter, 2005) because it is more difficult to apply such ideas in macro-food webs (insects, spiders, myriapods) where resource-based energy channelling is reticulated (Wolkovich, 2016; Potapov *et al*., 2021b).

Another set of studies diagnosed soil food-web structure and functioning using the abundance distribution of body size classes of soil biota from bacteria to earthworms, the idea mainly developed by Mulder and colleagues (Mulder, 2006; Mulder, den Hollander, & Hendriks, 2008; Mulder & Elser, 2009). The core idea of this ‘allometric’ approach is that abundance – body mass can serve as an indicator of environmental changes and is also linked to ecosystem functions, performed by different size classes (Petchey & Belgrano, 2010; Turnbull *et al*., 2014). The link between size spectrum and food-web structure is based on the assumption of a linear correlation between body size and trophic level across the food-web. However, this correlation is weak and multidirectional in soil (Potapov *et al*., 2021b). The size spectrum approach is also simplistic because it does not account for traits other than body size, such as food resource preferences and the spatial distribution of soil organisms.

The importance of the spatial distribution of energy fluxes in soil food webs was emphasized on the microscale, e.g. rhizosphere processes, on the macroscale, e.g. below-above ground energy transfer by mobile fauna (Scheu, 2001; Wardle *et al*., 2004) and for the horizontal patchiness of soil communities (Ettema & Wardle, 2002). For example, soil food-web structure vary with soil depth due to the differences in vertical distribution of different functional groups of soil fauna (Berg & Bengtsson, 2007). However, there is no systematic study of the spatial organization of energy channelling in soil food webs beyond the microscale.

Three, mostly independent, research directions are suggested by the literature overview above. These three correspond to three dimensions of soil food-web structure: (1) the resource-based energy channelling, (2) the body size distribution and (3) the spatial organization of trophic interactions in soil. Jointly, these structural dimensions are able to describe various aspects of functioning of soil food webs and their role in terrestrial ecosystems. But, so far, soil food-web reconstructions were focused on a single food-web dimension. Moreover, trophic interactions in soil food webs are generally reconstructed based on uncertain knowledge or on traditional assumptions about of what interactions occur. How the interactions are identified often lacks transparency and thus cannot be applied across different soil communities. Such reconstructions may, therefore, lack precision and this lack may affect the ecological conclusions drawn from them. The trophic interactions thus need to be validated against empirical *in situ* evidence, which has rarely been done before.

### (4) Revision of belowground food webs with novel tools

The methodological toolbox in soil trophic ecology is now much more diverse than it was some 20-30 years ago. Novel tools such as stable isotopes, fatty acids, and gut DNA analyses provide more realistic empirical descriptions of trophic links and food-web structure in cryptic belowground communities (Brose & Scheu, 2014; Potapov *et al*., 2021a). The spread of such novel tools has changed the understanding of soil food-web structure and functioning (Bradford, 2016). It has become evident that a major part of the energy fuelling soil food webs is root-derived (Ostle *et al*., 2007; Pollierer *et al*., 2007) and channelled through both bacterial and fungal pathways (Pollierer *et al*., 2012; de Vries & Caruso, 2016). It was also emphasized, that feeding across multiple energy channels is widespread for most of belowground consumers, including many microfaunal groups (Digel *et al*., 2014; Geisen, 2016; Wolkovich, 2016). At the same time, a range of feeding strategies was revealed in decomposer mesofauna groups, such as Collembola and Oribatida (Maraun *et al*., 2011; Potapov *et al*., 2016). The role of ectomycorrhizal mycelia has been challenged as a major food resource for soil fauna (Potapov & Tiunov, 2016; Bluhm *et al*., 2019), while soil autotrophic microorganisms were potentially overlooked one (Schmidt, Dyckmans, & Schrader, 2016; Seppey *et al*., 2017; Potapov, Korotkevich, & Tiunov, 2018). However, these findings are largely ignored in existing soil food-web reconstructions. Together with a number of experts in soil ecology, I attempted to synthesize classic knowledge with these recent findings by reviewing literature on the feeding habits of individual animal groups in a review that accompanies this paper (Potapov *et al*., [bioRxiv]). The conceptual paper presented here is based on this review and aims at developing a holistic approach to describe soil food webs across their resource-body size- and spatial dimensions and deliver a set of functional indicators, describing effects of consumers on ecosystem functioning and stability. In the following chapters I first revise generic food-web organization principles in relation to the soil system, then describe the multichannel food-web reconstruction approach and suggest functional indicators illustrating them with a hypothetical and an empirical example, and finally discuss the limitations of the approach, the main knowledge gaps and a way forward for the soil food-web research.

## II. Essential concepts in functional soil food-web research

### (1) Basal resources of soil food webs

Energy in food webs flows through consumer trophic chains, which may be clustered in energy channels based on a certain similarity. Resource-based energy channelling clusters trophic chains on the basis of the basal resources they use. This is probably the most common way to understand the structure and functioning of belowground food webs (Fig. 1A). However, the classification of basal resources and corresponding energy channels is often unclear. For example, the traditional distinction of root, bacterial and fungal energy channels (Hunt *et al*., 1987) is hardly applicable to macrofauna detritivores feeding mainly on litter and soil organic matter, and ignores autotrophic microorganisms. The general distinction between green and brown (i.e. grazing and detrital) channels (Moore *et al*., 2004) introduces ambiguity in the case of food chains based on root exudates and mycorrhizal fungi that are associated with living plant roots (i.e. green energy channel) being intimately interlinked with soil organic matter sequestration and decomposition (i.e. brown energy channel). Revision of these concepts should be the subject of a focused study introducing ontologies for reducing ambiguity and increasing the reproducibility of soil food-web research. In the present paper I understand basal resources as being the main organic pools at the base of the soil food-web that support soil consumers and that are associated with different ecosystem-level processes (Fig. 2). Different feeding adaptations are needed to consume different basal resources and by feeding on different resources consumers affect different ecosystem processes, including transformation, translocation and decomposition of organic matter, nutrient mineralisation, plant growth, microbial dispersal and others (Briones, 2014; Potapov *et al*., [bioRxiv]).

**Figure 1.**
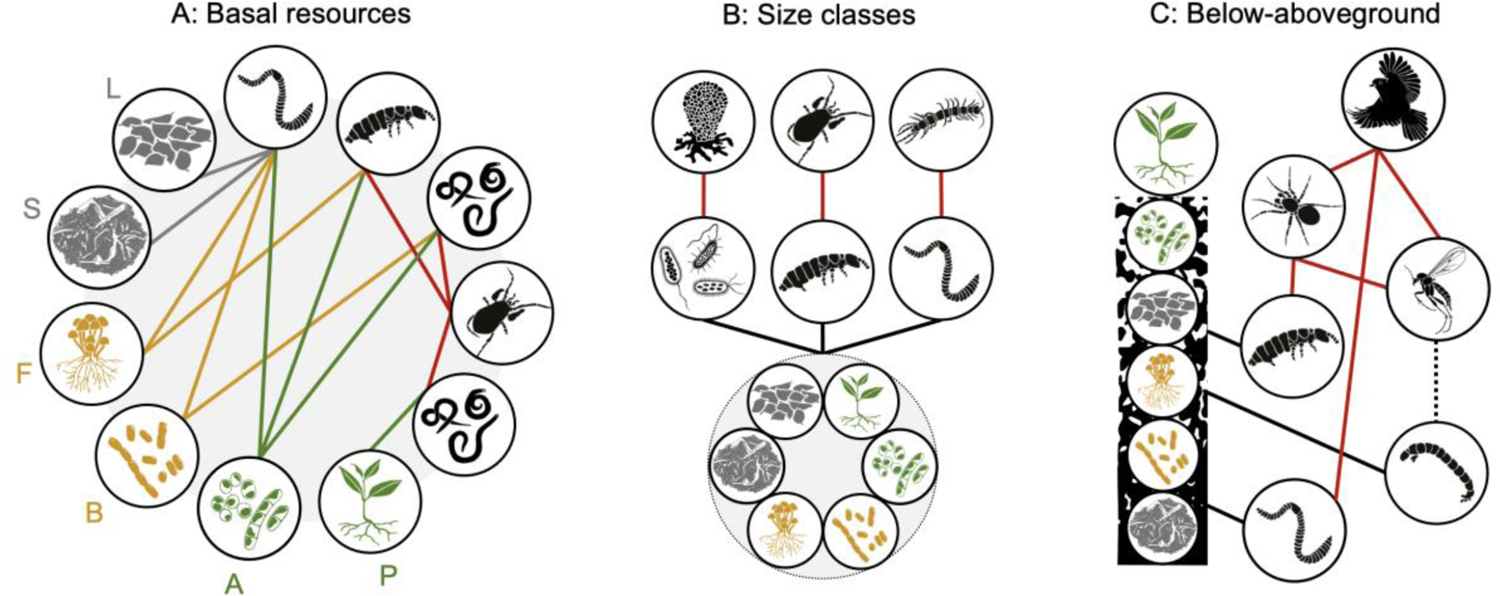
Structural facets of energy channelling in soil food webs. (**A**) Energy channels based on different resources have different turnover rate and control different ecosystem-level processes such as herbivory, decomposition and nutrient cycling. Brown channel unite detrital (grey), fungal and bacterial channels (dark yellow); green channel is based on living autotrophic organisms (green); predators couple different resource-based channels (red). For resource abbreviations refer to Fig. 2. (**B**) Different size classes of soil consumers impact different ecosystem functions and are controlled by different environmental factors. Energy from all resources is channelled in parallel via several size-based energy channels, each coupled by different predatory groups. (**C**) Consumers in the soil rely on spatially structured basal resources, translocate organic matter vertically and horizontally, subsidizing aboveground predators with prey biomass via vertical movements of soil fauna and winged insects that developed in the soil.

**Figure 2.**
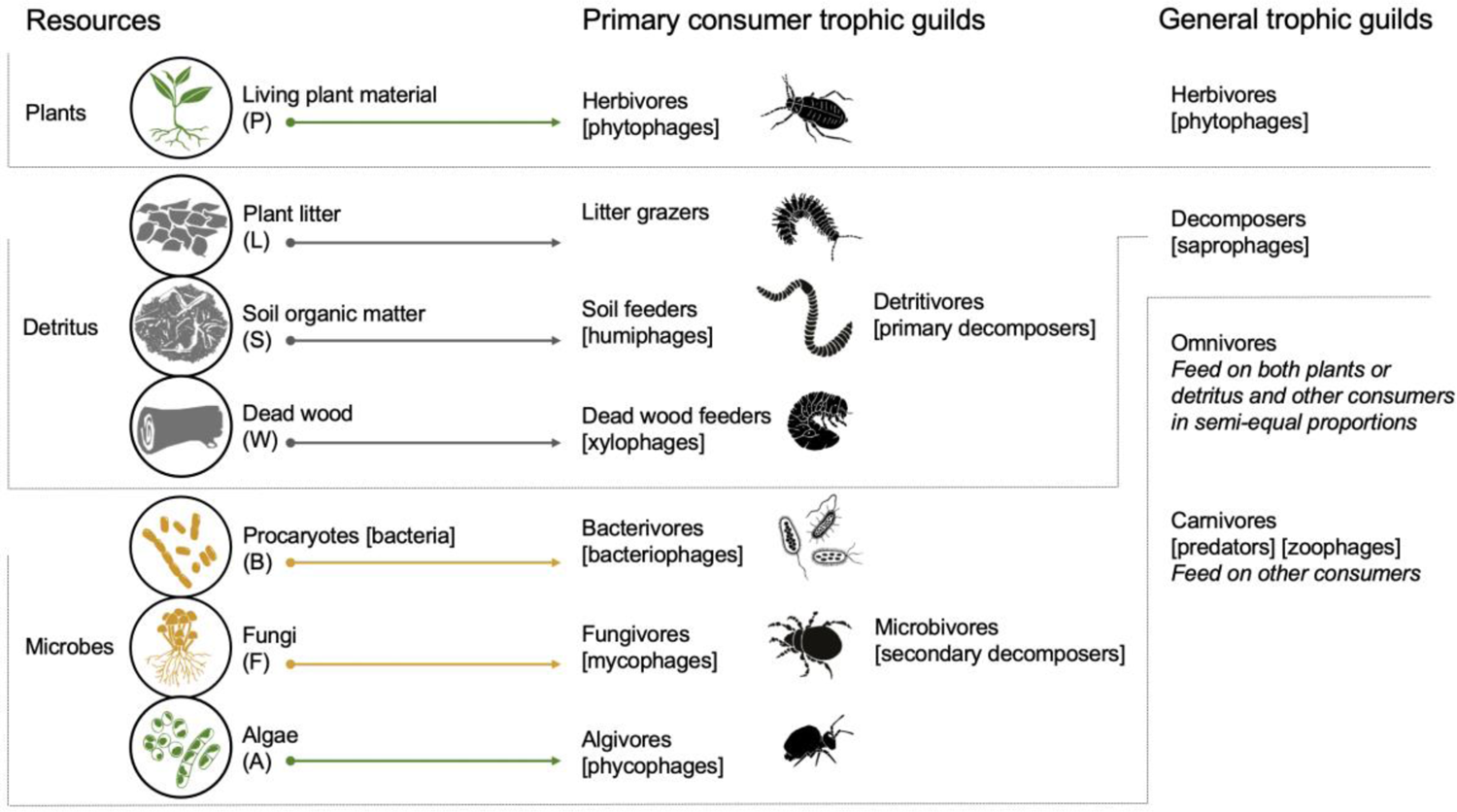
Basal resources and corresponding consumer trophic guilds in soil food webs. Animals and protists feeding on both detritus and microorganisms form a general guild of ‘decomposers’ that affect decomposition via food consumption. Decomposer prokaryotes and fungi, i.e. ‘saprotrophs’, are considered as resources in the present framework. Abbreviations for resources are given in brackets; synonyms are given in square brackets. Colours highlight brown (grey and dark yellow) and green energy channels (green). Dissolved organic matter is assumed to be used primarily by procaryotes and fungi and thus is not explicitly considered here. Summarised from (Swift *et al*., 1979; Striganova, 1980; Hunt *et al*., 1987; Potapov *et al*., [bioRxiv]).

### (2) Resource stoichiometry and assimilation efficiency

Basal food resources of soil food webs vary greatly in their elemental proportions and profitability for consumers. Assimilation efficiency, i.e. the proportion of ingested food that is assimilated by consumer, is different between resources such as living plants, detritus, microorganisms and animal tissues (Jochum *et al*., 2017; Lang *et al*., 2017). Detritivores have to eat more food to keep stoichiometric ratios of C:N:P in their bodies (Pokarzhevskii *et al*., 2003; Jochum *et al*., 2017) and thus consume a larger volume of food than e.g. predators do.

Detritivores with a low assimilation efficiency exhibit the largest effects on their environment via feeding activities, as exemplified by earthworms that consume hundreds of tons of soil per hectare per year (Lavelle & Martin, 1992). Thus, assimilation efficiency is one of the important parameters for quantifying ecosystem-level effects of resource-consumer interactions (see below “Metabolic ecology and energy flux approach”). Assimilation efficiency is predicted well by using, for example, the nitrogen concentration of the food resources (Jochum *et al*., 2017). Accounting for nitrogen concentrations or C to N ratios in resources and consumers is thus a promising approach for predicting interaction strengths in soil food webs (Buchkowski & Lindo, 2020).

### (3) Trophic guilds and taxonomic groups

Food webs are often reconstructed based on ‘trophic species’ that represent groups of biological species that share a similar pool of resources and predators (Yodzis & Winemiller, 1999; Luczkovich *et al*., 2003). In soil, such groups are traditionally termed trophic guilds and have a more functional focus, being linked to the exploitation of a specific basal resource in a specific way, and even having similar microhabitat preferences (Moore, Walter, & Hunt, 1988; Brussaard, 1998). To get correct estimates of food consumption, such groups should also share similar physiology and stoichiometry (Buchkowski & Lindo, 2020). In most reconstructions, pure trophic classification such as detritivores, bacterivores, fungivores, herbivores, carnivores and omnivores are mixed with high-rank taxonomic classifications (Hunt *et al*., 1987; de Vries *et al*., 2013; Gongalsky *et al*., 2021). This is justified, not only because taxonomic identification is the basis of food-web research, but also because trophic niches in soil fauna can, to a large extent, be predicted using phylogenetic (taxonomic) relationships among groups (Cardoso *et al*., 2011; Potapov *et al*., 2016; Potapov, Scheu, & Tiunov, 2019c). A hybrid taxonomic and guild approach (Brousseau, Gravel, & Handa, 2018; Laigle *et al*., 2018) also allows consideration of a number of phylogenetically conserved traits such as physiology, stoichiometry, and reproductive and defence strategies. Even though there are several reviews of the commonly used trophic guilds and functional groups (e.g. Moore *et al*., 1988; Brussaard, 1998; Briones, 2014), no comprehensive trophic guild classification across size classes in soil has until now been compiled. Nor has a common vocabulary across taxa been clearly defined. For the needs of the present study, I listed commonly used trophic classifications corresponding to the basal resources in Fig. 2. In the reconstruction below, I rely on the multifunctional classification compiled in the accompanying review (Potapov *et al*., [bioRxiv])

### (4) Size classes of soil consumers

Consumers, from protists to large invertebrate, may span from few micrometres to dozens of centimetres in body length and over twelve orders of magnitude in body mass in a single soil community (Potapov *et al*., 2021b). Body size is a very general trait that affects a number of organism characteristics including metabolism, growth rate and trophic interaction partners among others (Brown *et al*., 2004; Woodward *et al*., 2005). Different size classes in soil inhabit different environments (water, air pores and holes, or bulk soil), have different mobility restrictions and vertical stratification, exhibit different degree of trophic specialization, and vary in their engineering roles (Fig. 3) (Scheu & Setälä, 2002; Wardle, 2002; Briones, 2014; Erktan, Or, & Scheu, 2020). However, body size is not related to the trophic level across the food-web since top predators are present in different size classes (Fig. 1B) (Potapov *et al*., 2019a, 2021b). The conventional classification into micro-, meso- and macrofauna is based primarily on body width, since it is the main characteristic that restricts movement abilities of organisms in the soil (Swift *et al*., 1979). However, for food-web analysis, living body mass is very important since it provides information on which prey a predator is able to handle (Cohen *et al*., 1993). The body mass perspective results in elongated animals such as nematodes, myriapods and oligochaetes being assigned to larger size classes than those based on body width and in smaller size classes than those based on body length (Fig. 2) (Potapov *et al*., 2021b). Non-linear variations in trophic level with body mass suggest that small-sized soil-dwelling microarthropods are involved in micro-food webs together with microfauna, as depicted also in traditional soil food-web models (Hunt *et al*., 1987; Potapov *et al*., 2021b). By contrast, large microarthropods that live mostly in fresh litter and on the ground surface, are involved in macro-food webs (Potapov *et al*., 2021b). Describing soil communities using the size spectrum approach has the further advantage of correctly evaluating the food-web roles and ecosystem impacts of juvenile organisms (Potapov *et al*., 2021b; Gongalsky, 2021). Since different size classes of soil consumers impact different ecosystem functions and are controlled by different environmental factors, the size spectrum is an integrative indicator for the soil community (Mulder, 2006).

**Figure 3.**
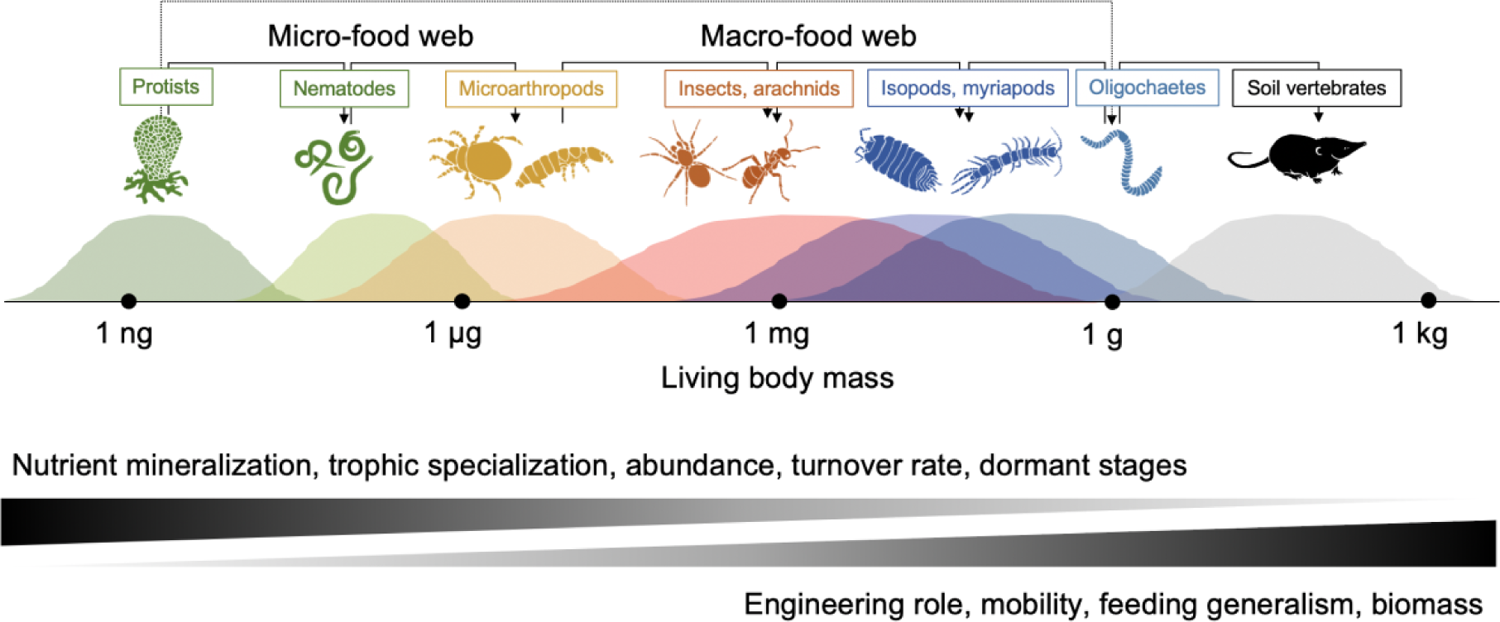
Body mass spectrum of consumers in soil. Well established soil communities embrace consumers spanning over twelve orders of magnitude in body mass. Small-sized consumers have a high turnover rate and affect nutrient cycling via microbial grazing. Large-sized consumers have a high biomass and play important engineering roles in soil via transformation and translocation of organic matter. Predator-prey trophic interactions (black arrows) occur predominantly among organisms of similar size, with micro-food webs being partially disconnected and consumed by macrodetritivores as a whole (protists-oligochaetes dashed arrow). Summarised from (Lavelle, 1996; Scheu & Setälä, 2002; Pokarzhevskii *et al*., 2003; Erktan *et al*., 2020; Potapov *et al*., 2021b).

### (5) Predator-prey interactions, mass ratios and traits

Generalist feeding is a common feature in soil food webs and is especially evident in predatory groups (Scheu & Setälä, 2002; Digel *et al*., 2014). However, generalist feeding greatly hinders the systematic occurrence of species-specific interactions in soil. Such interactions are rare because communication (whether chemical or in other forms) between animals in the soil is difficult and because community composition is very variable across space. When an empirical assessment of trophic interactions is not feasible, trophic interactions are reconstructed based on expert knowledge and existing evidence in the literature (Hunt *et al*., 1987; Digel *et al*., 2014). Generalist feeding, however, makes realistic the assumption that most of the possible interactions actually exist. Predator-prey mass ratios (PPMRs) can be used to define interactions that are possible physically (and energetically profitable) (Brose *et al*., 2008) (Fig. 4A). Body masses alone correctly predict more than 50% of trophic interactions across size classes of consumers in marine and aboveground food webs where trophic interactions are typically more specialized (‘allometric’ models, Petchey *et al*., 2008). The few PPMR estimates that exist for soil predators suggest that the optimum varies around 100, i.e. the predator is approximately 100 times heavier than its optimum prey (Brose *et al*., 2008). However, purely allometric models have a large uncertainty when being tested against empirical data on soil macropredators (Eitzinger *et al*., 2018). Indeed, predators may also feed on prey much smaller, or handle prey of comparable size, depending on specific predator and prey traits (Fig. 4B) (Brose *et al*., 2019). In soil communities, key traits that may modify PPMRs, presence and intensity of predator-prey interactions, are often attributed to certain taxonomic groups and include hunting adaptations and ingestion mechanisms of predators, as well as protective metabolites, physical structures and behaviour of prey (Table 1) (Brousseau *et al*., 2018; Laigle *et al*., 2018). For example, strongly sclerotized groups such as oribatid mites has been shown to be little attacked by predators (Peschel *et al*., 2006). However, the effectiveness of different protection mechanisms against different predators has not been systematically studied in soil. Together with body mass, these traits are expected to provide more realistic reconstructions of trophic interactions in generalist soil food webs.

**Figure 4.**
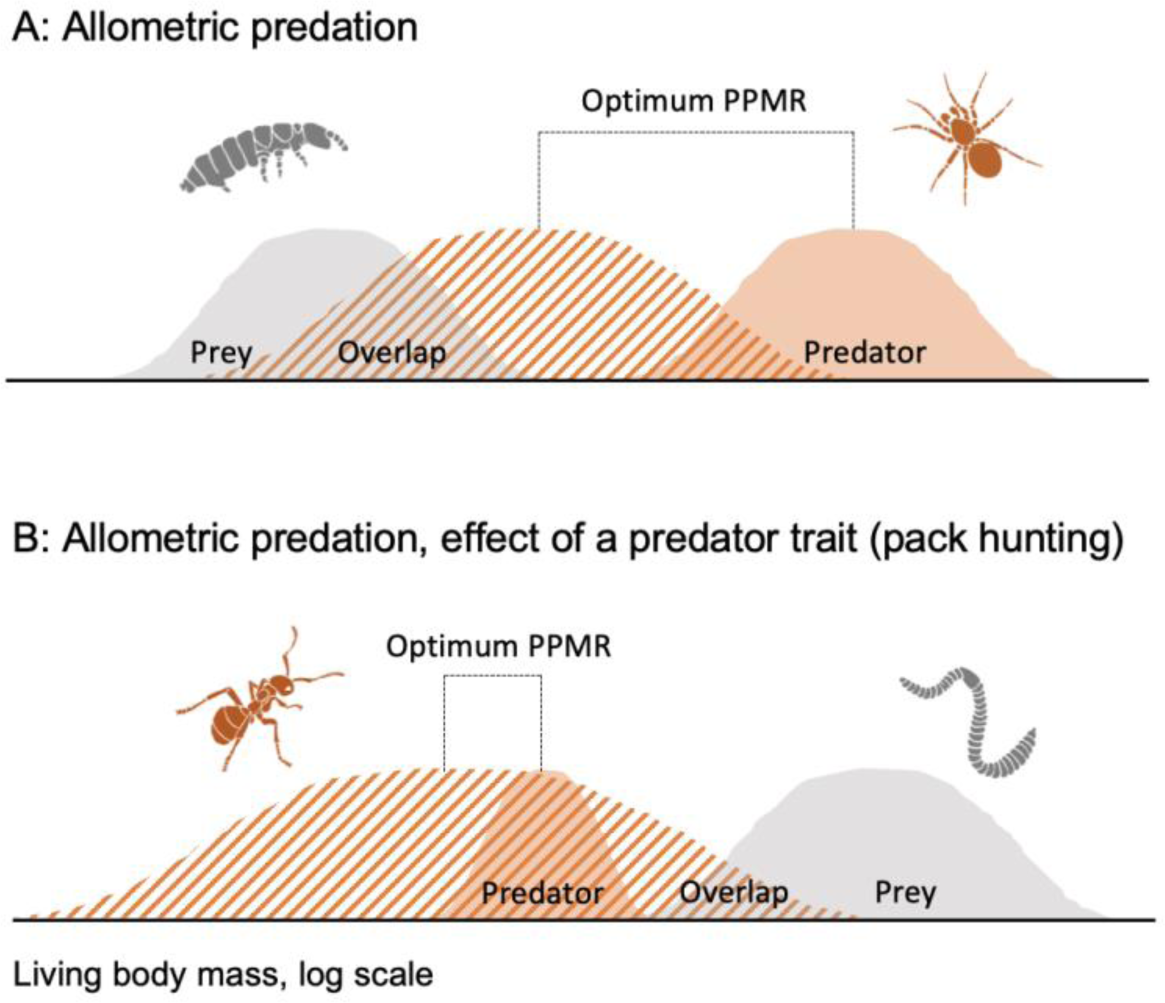
Feasible predator-prey interactions depend on body mass ratios. Small prey has a low handling time, but also is less energetically profitable than large prey, shaping an ‘optimum’ predator-prey mass ratio distribution (PPMR) (Brose *et al*., 2008). (**A**) Optimum PPMR together with population body mass distribution of a predator (dark orange-filled distribution) and a prey (grey-filled distribution) can be used to predict interaction strength between them (lined overlap with grey distribution). (**B**) Specific traits of predator and prey may modify PPMR and interaction strength. Despite ants being smaller than earthworms, pack hunting and venom shifts and widens the PPMR distribution, making the predation feasible.

**Table 1.**
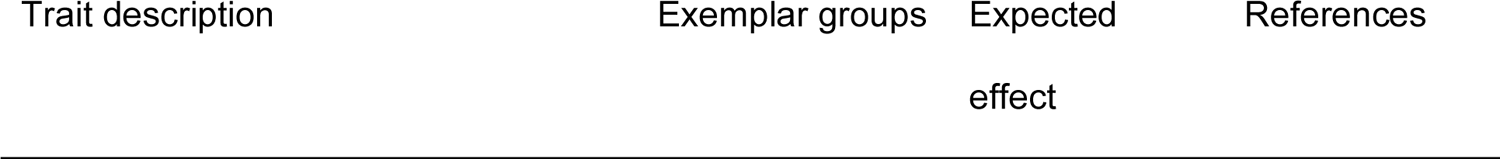
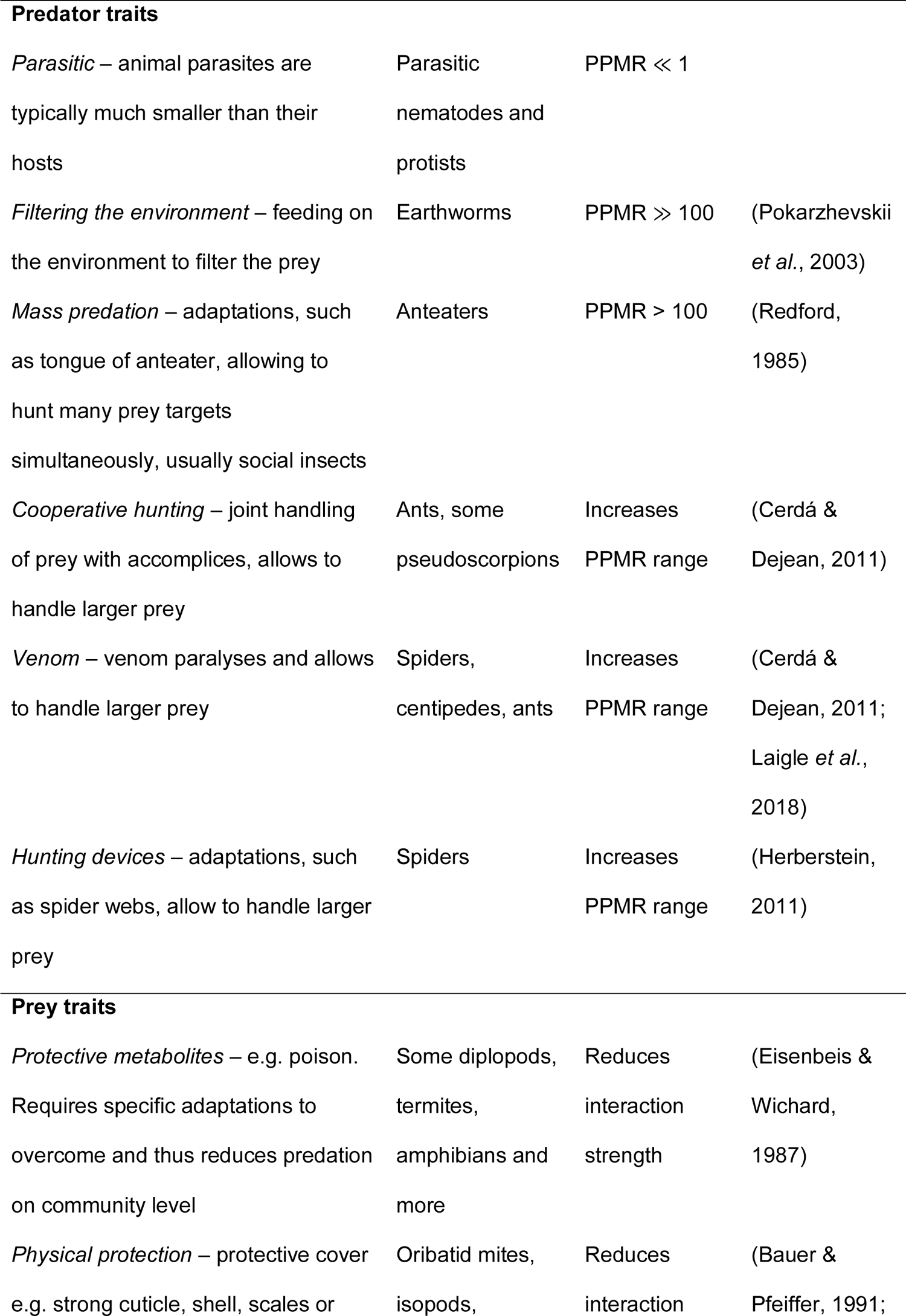
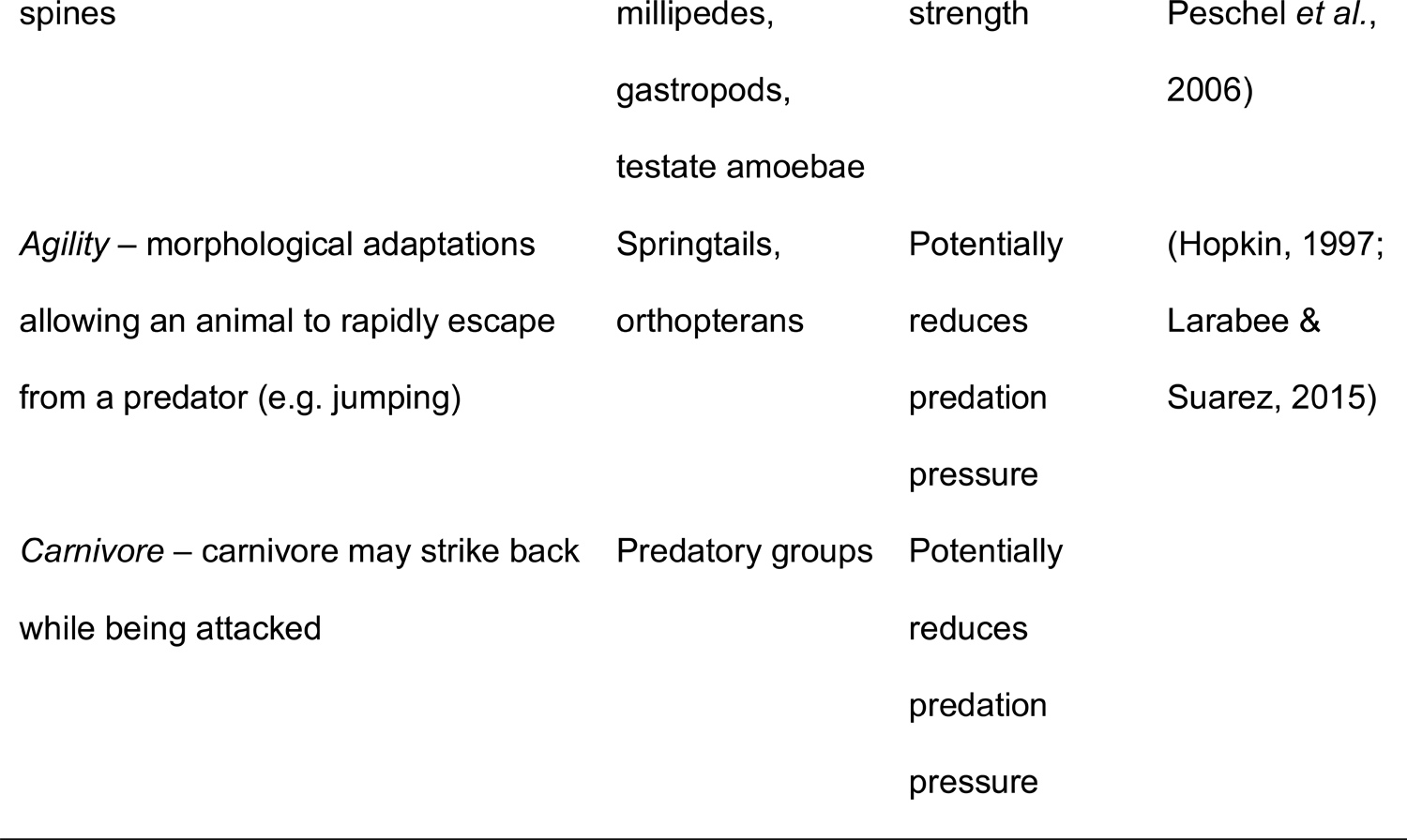
Predator and prey traits, modifying interaction strength and PPMRs. Numbers that are shown in the ‘expected effects’ column are general theoretical expectations and not strict rules.

### (6) Vertical stratification of soil food webs

Soil is a stratified environment so that there are divergent evolutionary pressures on fauna living on the surface, and those in the mineral soil (Ghilarov, 1949). These divergent pressures creates vertical stratification of forms and functions in soil communities (Ellers *et al*., 2018). Mobile ‘epigeic’ groups of arthropods inhabit the surfaces of fallen leaves, wood, stones, and bare soil; ‘hemiedaphic’ invertebrates inhabit coarse detritus, such as decomposing litter or wood; ‘endogeic’ invertebrates inhabit lower organic and mineral soil layers, including the rhizosphere. Classifications related to vertical stratification were developed in different soil taxa including e.g. earthworms (Bouché, 1977), springtails (Gisin, 1943) and gastropods (Ellers *et al*., 2018). Vertical stratification of taxonomic groups propagates to the corresponding vertical stratification in the structure and energy fluxes in soil food webs (Berg & Bengtsson, 2007; Okuzaki *et al*., 2009). Moreover, the spatial distribution of basal resources also structures energy channelling in soil food webs: fresh organic detritus and algae are more abundant in the surface layers, while soil organic matter and roots are more abundant in the mineral soil (Fig. 1C) (Ponge, 2000). Vertical stratification also provides information on the spatial niche differentiation among different functional groups that limits predator-prey interactions among groups that live in different layers. Many large fauna, however, move vertically through the soil profile during their development, or depending on the environmental conditions (Dowdy, 1944). This pattern is especially evident for holometabolous insects (flies, beetles) many of which have larval stages in the soil and flying adults (Ghilarov, 1949). But even true soil-dwelling invertebrates, such as earthworms, channel energy from soil organic matter ‘directly to the sky’ when they are eaten by birds (Fig. 1C). Detrital subsidy is probably indispensable for aboveground predators in virtually all terrestrial ecosystems, but its quantification and origin have seldom been studied (Scheu, 2001; Hyodo, Kohzu, & Tayasu, 2010; Hyodo *et al*., 2015).

### (7) Metabolic ecology and energy flux approach

The impacts of soil animals and protists on ecosystem functioning are, in most cases, linked to the consumption of other consumers, microorganisms, litter and soil and burrowing in search of food. Consumption rate, in turn, is defined primarily by the metabolic demands of an organism – the amount of energy it needs to sustain its life. The metabolic rate scales sublinearly with living body mass, varies with phylogenetic position of the consumer, and increases steadily with environmental temperature (Brown *et al*., 2004; Ehnes, Rall, & Brose, 2011). Metabolic rate accounts for the different metabolic demands of small and large organisms per unit of body mass and is thus a universally comparable measure of organism and population impacts on ecosystem functioning across size classes, superior to biomass or numeric abundance. Resource consumption rate depends primarily on the metabolic demands of an organism and on the efficiency with which it can assimilate its food resources (see above “Resource stoichiometry and assimilation efficiency”). In an energetically steady-state system (i.e. losses equal gains for each food-web node), consumption rate also depends on the position of a consumer in the food-web because lower trophic levels sustain higher trophic levels with energy. Consumption rate, after accounting for the assimilation efficiency and losses of energy to higher trophic levels in the food-web, represents the total energy flux out of all resource nodes to a consumer, which can be used as a measure of its ecosystem-level impact (Barnes *et al*., 2014, 2018). The energy flux approach was applied in traditional soil food-web models to quantify the contribution of soil consumers to nitrogen mineralisation (Hunt *et al*., 1987; de Ruiter *et al*., 1993). More recently, the approach was linked to biodiversity and expanded to more diverse ecosystem functions (Barnes *et al*., 2014). In that paper, ecosystem functions such as herbivory, decomposition and predation were inferred from the energy fluxes to corresponding trophic guilds of macroinvertebrates.

Widespread application of this approach to soil food webs, however, is hampered by the generalist feeding of soil animals and their poorly documented feeding preferences. Both these factors often make trophic guild assignment unrealistic. I think that we need more realistic reconstructions, incorporating different aspects of detritivory, widespread omnivory and multichannel feeding, and specific body size and spatial structures of soil food webs. In the chapters below, I build on the classification of soil consumers given in an accompanying review (Potapov *et al*., [bioRxiv]) to reconstruct soil food webs. I then use the energy flux approach, as implemented in the R package *fluxweb* (Gauzens *et al*., 2019), to calculate indicators of their functioning. The suggested multichannel reconstruction unites the resource, size and spatial dimensions of soil food webs and can be applied from a local to a global scale.

## III. Reconstruction of multichannel food webs

### (1) Food-web reconstruction

I propose a novel ‘multichannel’ approach of soil food-web reconstruction which predicts trophic interaction strengths in a given soil community using prior knowledge of species biology, basic food-web principles and the key traits of consumers. Multichannel reconstruction of soil food webs relies on trophic guilds as the network nodes that are distinguished based on multiple trait similarities. The assignment of traits to groups can be based on published, or directly measured, empirical data. The reconstruction of trophic interactions among groups is based on trait relationships that are extrapolated from existing experiments to generic rules. The approach thus produces a hypothetical food-web structure that describes reality as well as possible given current knowledge. The reconstruction approach is conceptually close to the multidimensional niche model of food-web reconstruction, which assumes that trophic interactions are formed and selected along several trait dimensions (Allesina, Alonso, & Pascual, 2008). Here, these dimensions are represented by phylogenetically-defined feeding preferences, body sizes, protection mechanisms and vertical stratification as well as other traits that are expected to modify PPMR or consumption rate (see “Essential concepts in functional belowground food-web research” above). The full list of trophic guilds and corresponding traits is compiled in the accompanying “Multifunctional trophic classification of belowground consumers from protists to vertebrates” (Potapov *et al*., [bioRxiv]) and shortlist is given in the Supplement (Table S1). The following main assumptions were used to reconstruct trophic links (step-by-step reconstruction is given in the Appendix S2):

1. There are phylogenetically-driven differences in feeding preferences for various basal resources and predation capability among soil animal taxa that define their feeding interactions (Laigle *et al*., 2018; Potapov *et al*., 2019c). These preferences were assigned based on the accompanying review (Potapov *et al*., [bioRxiv]) (Table S1).
2. Predator-prey interactions are primarily defined by the optimum PPMR. Typically, predator is larger than the prey, but certain predator traits (hunting devices and behaviour, parasitic lifestyle) can considerably modify the optimum PPMR (Fig. 4).
3. Strength of the trophic interaction between a predator and a prey defined by the overlap in their spatial niches related to vertical differentiation. The higher is the overlap, the stronger is the interaction (Table S1).
4. Strength of the trophic interaction between a predator and a prey can be considerably reduced due to the prey protective traits (Peschel *et al*., 2006) (Table 1).
5. Predation is density (biomass) dependent (Gauzens *et al*., 2019). Due to a higher encounter rate, predator will preferentially feed on prey that is locally abundant.

To illustrate the multichannel reconstruction, I selected groups from the list of trophic guilds of soil consumers that commonly co-exist in the soil food webs of temperate forests (Table S1). I followed assumptions 1-4 described above to produce an interaction matrix (adjacency matrix). Similar reconstruction approaches have been applied to invertebrate food webs (Digel *et al*., 2014; Hines *et al*., 2019). Here, however, I make the assumptions behind such reconstructions more complete and better reproducible for future studies. The reconstruction included soil consumers spanning from primary consumers to intraguild predators and from protists to vertebrates (Fig. 5). In this reconstruction, I assumed that biomasses of all nodes are equal and ignored different metabolic losses across nodes and thus the interaction strengths in this hypothetical example represent feeding preferences rather than energy fluxes, with the goal of illustrating the concept. By combining this reconstruction with empirical biomass data, it was possible to calculate energy fluxes across the food-web with the *fluxweb* package (Gauzens *et al*., 2019; Jochum *et al*., 2021).

**Figure 5.**
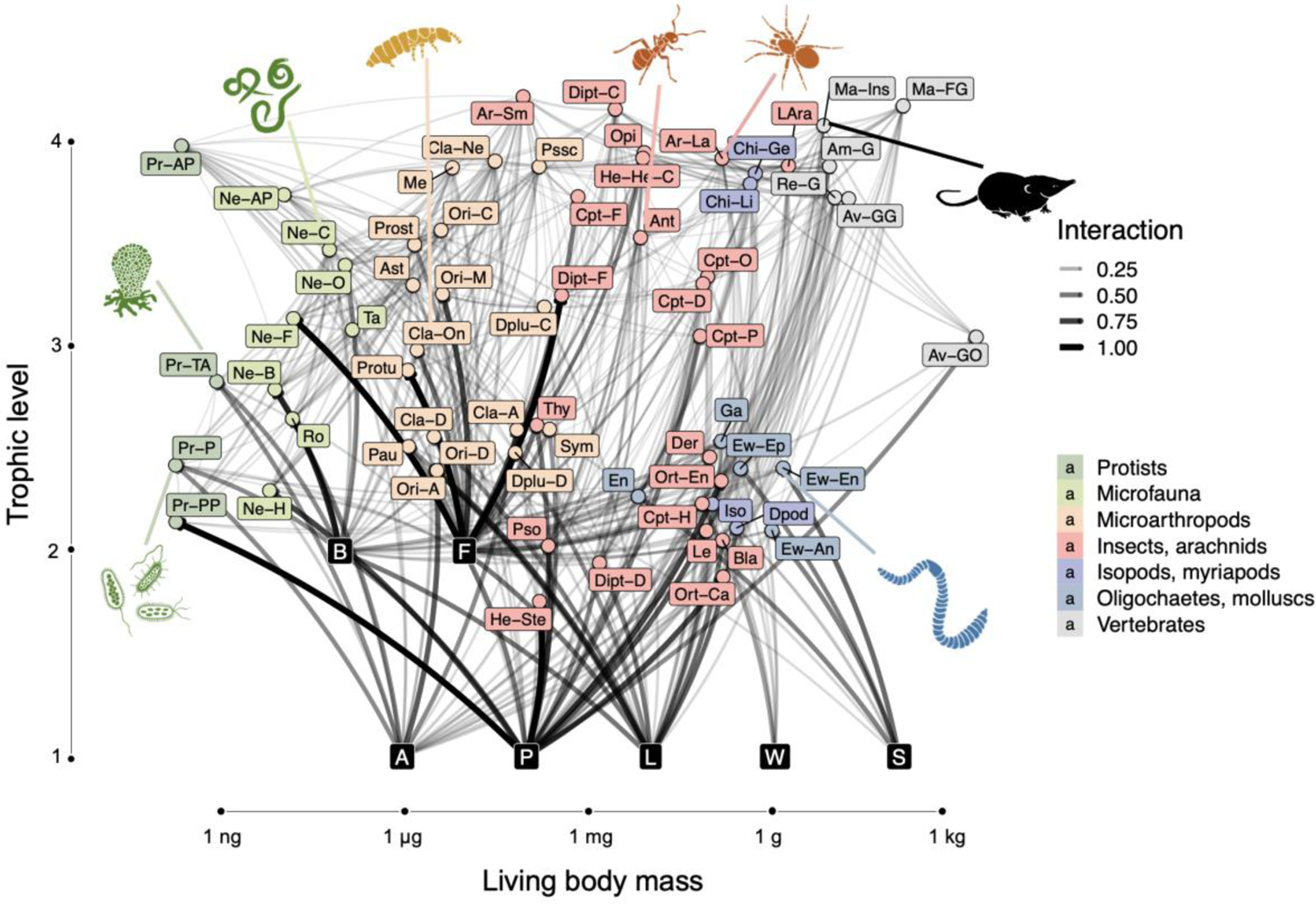
Reconstructed multichannel meta-food-web of temperate forests, from protists to vertebrates. The 66 consumer nodes are represented by trophic guilds and are arranged based on body mass (x axis) and trophic level (y axis). Node colours indicate broad quasi-taxonomic groups. Resources are shown with black square labels and positioned arbitrary along the body mass axis. Procaryotes and fungi are positioned at trophic level two as they receive energy from dead or living primary producers (trophic level one). Width and darkness of links indicates feeding preferences inferred from spatial, size, protection and feeding preference adjacency matrices (only interactions comprising >2% of the energy budget of the consumer node are shown). Biomass is assumed to be the same across all nodes and so biomass-dependent feeding preferences are not included. For resource abbreviations refer to Fig. 2; for trophic guild abbreviations refer to Table S1.

### (2) From interactions to functions

Once a food-web is reconstructed and the energy fluxes are calculated, there is a great opportunity to quantify different aspects of how consumers contribute to ecosystem functioning (see “Metabolic ecology and energy flux approach” above). In comparison with existing applications of the energy flux approach, I treated most of basal consumers as omnivores, having many basal resource – consumer links and being involved in different resource-based energy channels (Fig. 6A). Omnivory of basal consumers represents the multichannel feeding widespread in soil decomposers (Brose & Scheu, 2014; Wolkovich, 2016) and actually prevalent in most natural food webs (Thompson *et al*., 2007; Wolkovich *et al*., 2014). Ecosystem functions can be still inferred from such complex food webs by summing up individual fluxes: e.g. summing up all outgoing fluxes from plants to other food-web nodes reflects absolute herbivory (Barnes *et al*., 2020) (Fig. 6A “Ecosystem functions”). Focusing on resource-based energy fluxes, it is possible to quantify bacterial versus fungal energy channelling, a measure suggested to indicate food-web stability (Moore *et al*., 2005) and nitrogen mineralisation (de Vries *et al*., 2013). Resource-based channels are not linked to the trophic functions alone. For example, the energy flux from litter represents consumption of litter by the consumer community, and thus is related to litter decomposition, transformation and translocation. The energy flux from soil organic matter similarly represents the consumption of soil and thus soil organic matter transformation and translocation, being linked to biopedoturbation and the modification of soil structure (Fig. 6B). Such inferences from energy fluxes about effects that are not purely trophic are justified particularly in soil where the habitat and the food are tightly interlinked (Fujii, Berg, & Cornelissen, 2020).

**Figure 6.**
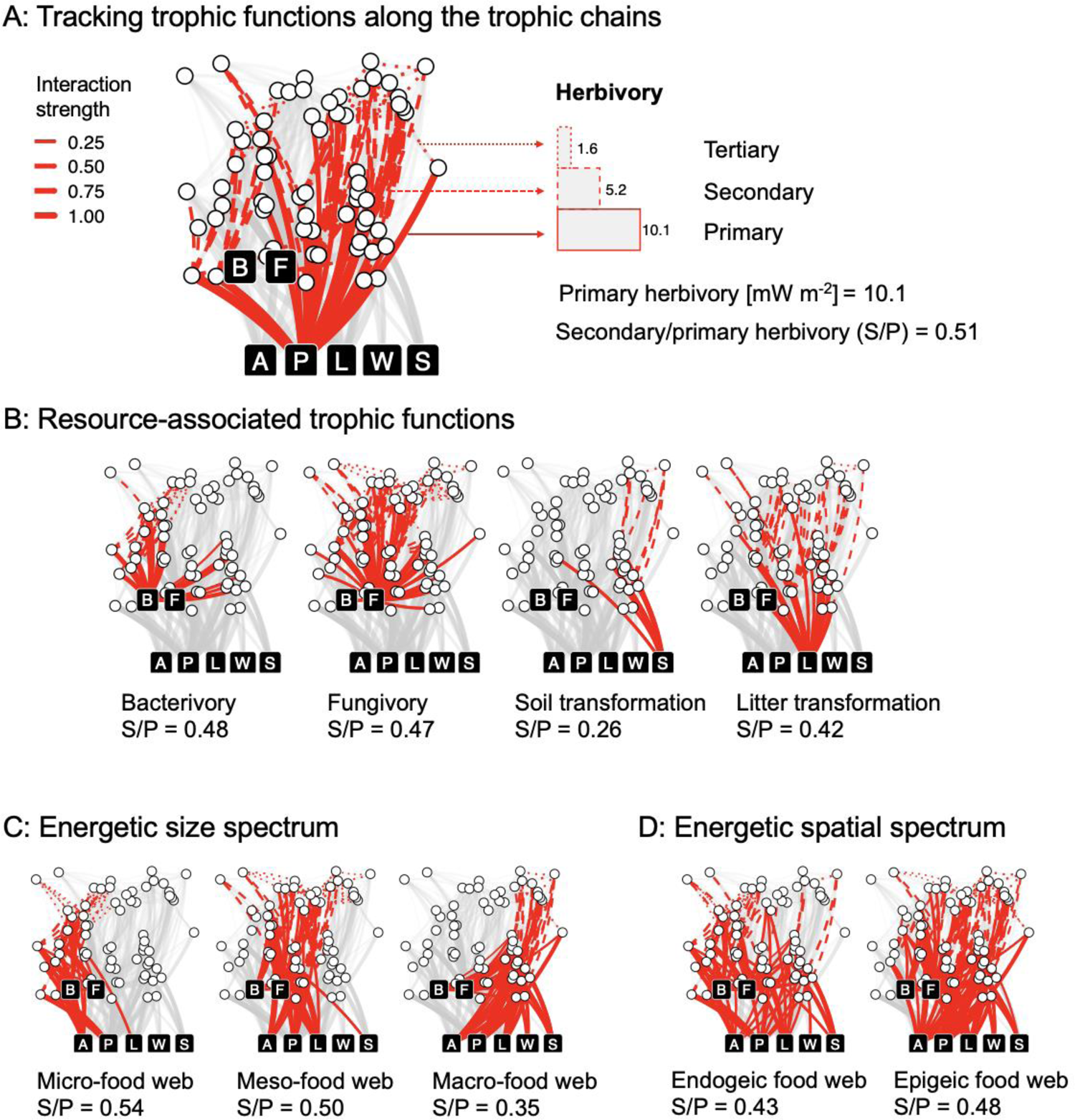
Inferring ecosystem functioning based on energy fluxes in soil food-web. The sum of outgoing energy fluxes from a specific food resource represents the total consumption of this resource and is thus related to corresponding ecosystem function(s). Interactions of the first (solid lines), second (dashed) and third (dotted) trophic levels indicate the proportion of energy that is channelled to the next trophic level through a specific channel. The ratio of primary to secondary energy flux (S/P) is related to the energy transfer efficiency and top-down control in the corresponding energy channel, or the entire food-web. For example, the sum of energy fluxes from plants to plant consumers is related to the ecosystem-level herbivory (primary herbivory) (**A, B**). Channelling of energy through micro-, meso- and macro-food-web compartments is related to a number of ecosystem-level processes that are driven by different size classes of soil consumers (see Fig. 2) (**C**). Channelling of energy through endogeic or epigeic food-web compartment is related to detrital subsidy and above-belowground interactions (**D**). Line thickness is proportional to interaction strength. A full version of the analysed network is represented in Fig. 5.

To illustrate the size aspect of food-web compartmentalisation, I classified all consumers linked to basal resources into three size classes: micro (protists and microfauna less than 0.3 µg in living body mass), meso (mesofauna and small macrofauna from 0.3 µg to 10 mg) and macro (macrofauna more than 10 mg). By tracking outgoing fluxes from the basal consumers of different size, I was also able to show remarkable distinction between the size-based channels in predators (Fig. 6C). In my reconstruction this distinction arises from the assumption of allometric trophic interactions, however, it confirms existing empirical data and is therefore probably realistic (Potapov *et al*., 2019a, 2021b). The reconstruction of size-based channels illustrates that basal resources are exploited by different size classes in different proportions and feeding on multiple resources is more pronounced in large primary consumers (Fig. 3). The distribution of energy among different size classes is related to ecosystem functioning and can be used as an integrative functional descriptor of a food-web, i.e. “energetic size spectrum”. It better reflects ecosystem functions of consumers than size spectrum approaches based on biomass or community metabolism (Mulder *et al*., 2011; Ehnes *et al*., 2014) because it reflects multitrophic energy fluxes and thus consumption rates. Size-based energy channelling could be also combined with resource-based energy channelling e.g. to unravel the contribution to resource-related functions of different food-web size compartments.

To illustrate the spatial aspect of food-web compartmentalisation, I classified all consumers linked to basal resources into endogeic (living in soil and lower litter layer) and epigeic (living in the fresh litter and on its surface). Some predator nodes specialized on one of these channels although many were linked to both endogeic and epigeic channels (Fig. 6D). The differentiation between endogeic and epigeic channels was partly related to the body size classes since many large macrofauna predators are surface-dwelling. It was also partly related to resource use due to different resource availability in different layers. The energy flux through the endogeic channel is expected to be related to soil structure modification and rhizosphere processes but the energy flux through the epigeic channel is expected to be related to the detrital subsidy available for aboveground consumers (Hyodo *et al*., 2015). Soil food webs with high energy flux through the epigeic channel are expected to support a higher biomass and diversity of aboveground consumers (Potapov *et al*., 2021b). Classifying energy fluxes according to resource, size and spatial perspectives allowing to ask more precise questions related to the food-web functioning, e.g. which size class in which soil layer is responsible the most for processing of a certain resource?

### (3) Describing the trophic hierarchy of energy channels

Each energy channel relies on basal resources or consumers and can be tracked to higher trophic levels. For each predator, the contribution of different basal resources and pathways to these resources can be described and quantified. For each channel, the amount of energy that reaches higher trophic levels can be estimated. Summing up predator-prey energy fluxes allows the quantification of “secondary” and “tertiary” trophic functions, such as predation, intraguild predation and parasitism (Barnes *et al*., 2014; Potapov *et al*., 2019b). These functions can be related to the top-down control of the entire food-web, a specific energy channel, or a specific consumer. This can be quantitatively assessed by calculating the ratio of the outgoing energy flux from a food-web node to the biomass of the node (Barnes *et al*., 2020) or by calculating the ratio of the energy fluxes at the bottom and at the top of the food-web. Such “energy flux pyramids” of primary, secondary and third-level trophic interactions can vary substantially not only across food webs, but also within a food-web across different energy channels (Fig. 6A-D). Even though my reconstruction did not account for biomasses, I observed several expected patterns in the ratios of secondary to primary energy fluxes (S/P) based solely on feeding preferences. For example, top-down control was higher in micro- and meso-than in macro-food webs (Potapov *et al*., 2019b, 2021b) and it was higher for plant-based than for soil-based energy channels (Fig. 6A,B,C). A number of more specific research questions related to top-down or bottom-up controls can be addressed with the energy flux approach, depending on the completeness of the food-web reconstruction (Fig. 7).

**Figure 7.**
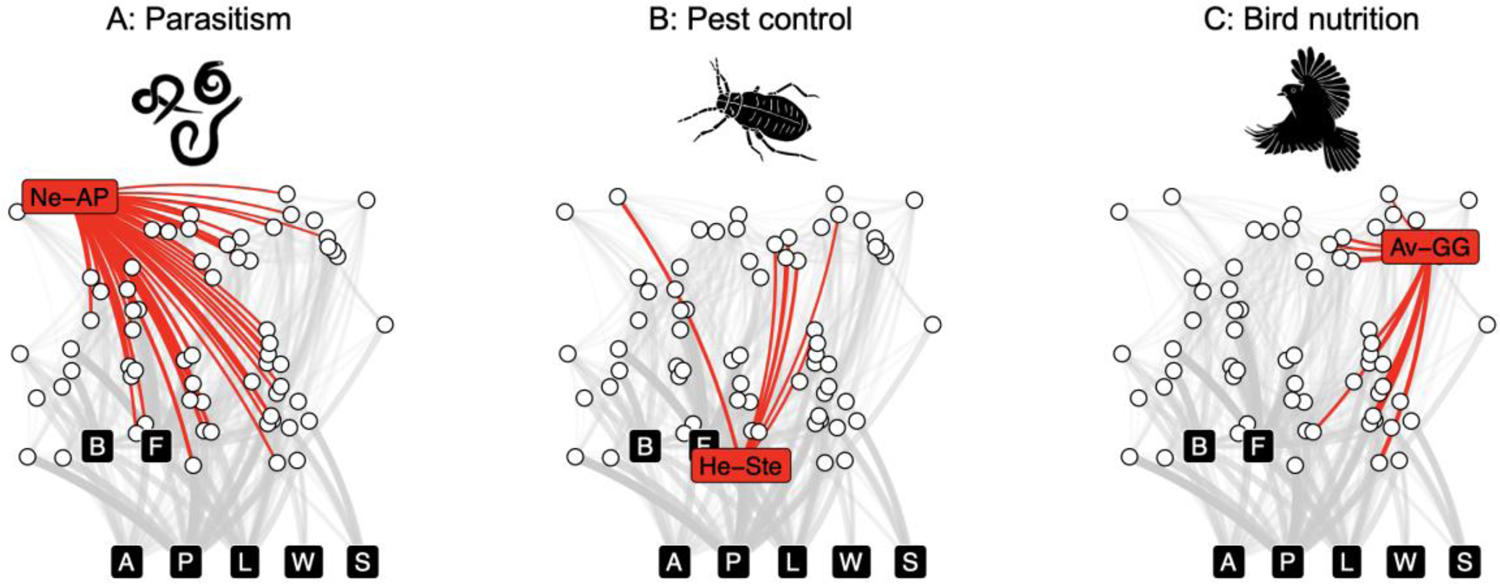
Assessing specific food-web processes with energy fluxes. The sum of energy fluxes from hosts to parasites can be used to assess community-level parasitism (**A**). The sum of outgoing energy fluxes from a pest can be used to quantify its top-down control (Barnes *et al*., 2020) and identify potential key biocontrol agents (**B**). The sum of incoming energy fluxes to a bird (or birds) can be used to identify potential animal groups and basal resources that play the most important role in its nutrition (**C**).

### (4) Assessing multifunctionality and energetic inequality

Ecosystem functioning is inherently multidimensional, which has fuelled a spectrum of studies assessing ecosystems by multiple functions simultaneously, i.e. ecosystem ‘multifunctionality’ (Wagg *et al*., 2014; Manning *et al*., 2018; Grass *et al*., 2020). Total energy flux in the food-web has been suggested as a proxy for ecosystem multifunctionality, since it comprises the sum of individual trophic functions. However, if a single trophic function predominates, total energy flux does not reflect multiple functions. In that case, approaches that consider all functions to be equally important are preferable (Potapov *et al*., 2019b). ‘Total flux’ and ‘average flux’ approaches resemble the ‘summing up’, ‘averaging’ and ‘threshold’ approaches commonly used to assess multifunctionality (Manning *et al*., 2018). By treating individual resource-size- and spatial energy channels as functions, it is possible to calculate different multifunctionality indices of soil food webs using the multichannel reconstruction (Table 2).

**Table 2.**
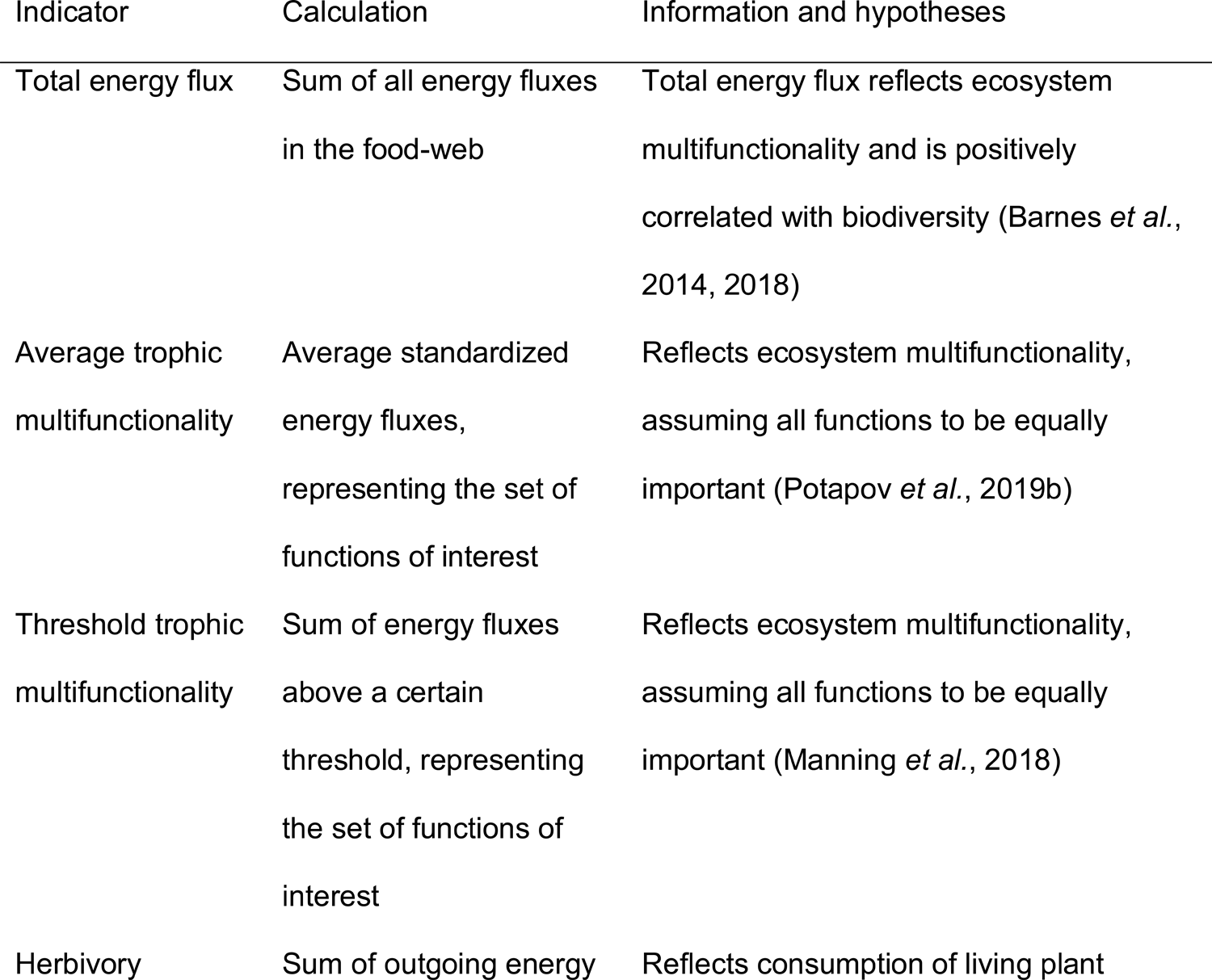
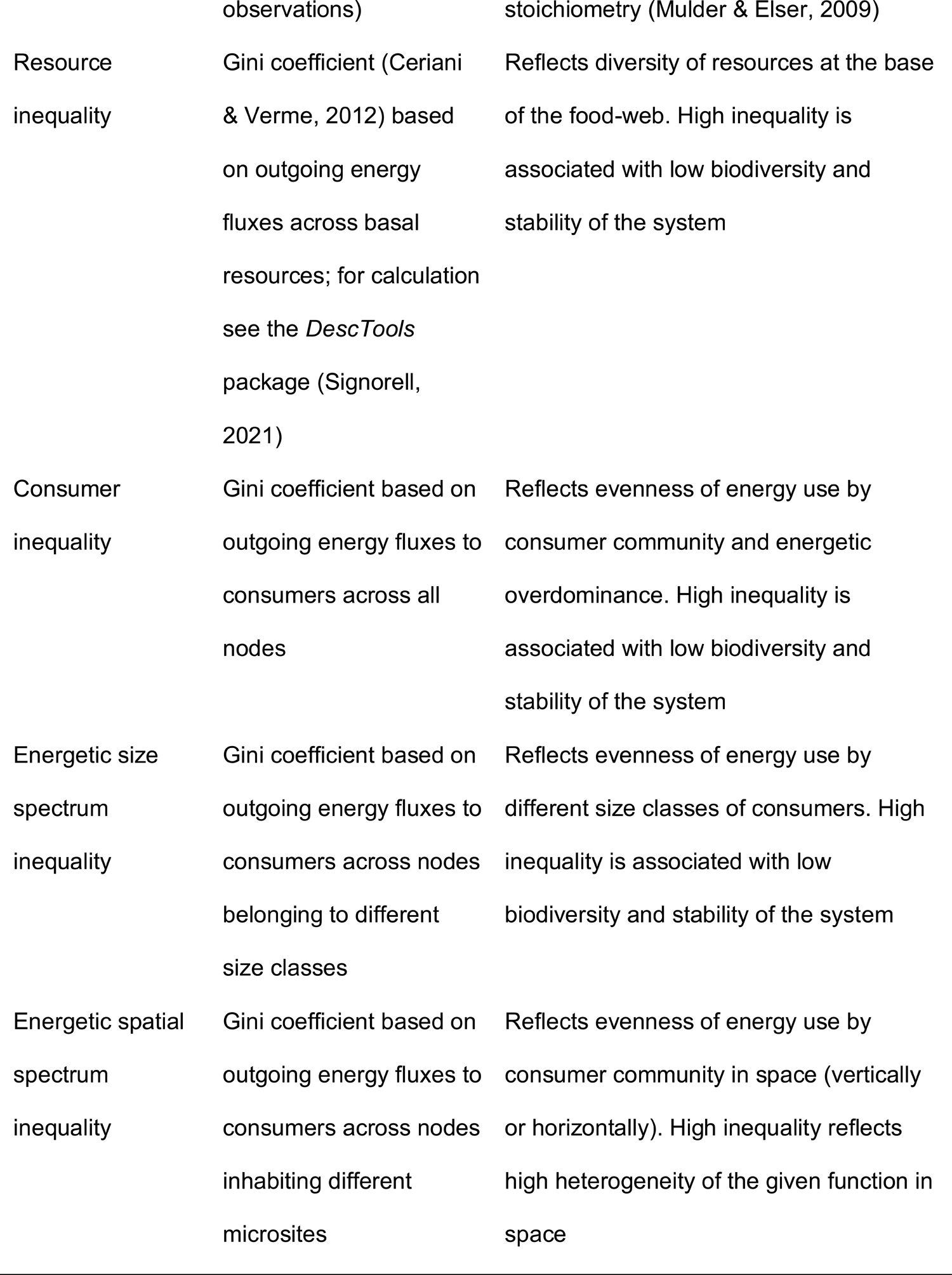
Functional indicators based on energy fluxes in the multichannel food-web reconstruction. List includes general indicators linked to the food-web functioning and stability, being non-exhaustive.

Another key aspect of the food-web is stability. The development of the resource-based energy channelling paradigm in soil food webs allowed the formulation of the concept of ecosystem stability as being driven by the balance between the fast (e.g. bacterial) and the slow (e.g. fungal) energy channels (Moore *et al*., 2005; Rooney *et al*., 2006). I think that this vision can be extended, for soil organisms, beyond ‘bacterial versus fungal’ energy channelling and include other resource and also body size energy channels (Potapov *et al*., 2021b). My hypothesis is that ecosystem stability and multifunctionality are both linked to the balance across different energy channels, decreasing in food webs with large energetic imbalances. Such imbalances can be observed across resources (e.g. bacteria-dominated systems), size spectrum (e.g. earthworm-dominated systems), spatial distribution (e.g. systematic ground surface disturbance) and trophic levels (e.g. systems with overdominance of primary consumers). Inequality could be quantified e.g. with Gini coefficients, widely used in social sciences to quantify income inequality (Table 2) (Ceriani & Verme, 2012).

### (5) Case study and comparison with traditional reconstructions

To illustrate the multichannel reconstruction with an empirical example, I re-analysed data on nematodes, mesofauna and macrofauna collected from rainforests and oil palm plantations in Sumatra, Indonesia (Krashevska, 2019; Potapov *et al*., 2019b). I used the empirical data on abundance and body masses together with trophic guilds (Table S1) to reconstruct the soil food webs. To demonstrate the significance of the reconstruction approach, I applied in parallel multichannel reconstruction (Fig. 8B) and traditional soil food-web reconstruction that assumes differentiated resource-based channelling in soil food webs (Fig. 8A) (Hunt *et al*., 1987; de Ruiter *et al*., 1993; Moore *et al*., 2005). Energy fluxes were assessed in both reconstructions with the *fluxweb* package (Gauzens *et al*., 2019) and used to calculate a set of functional indicators. Various technical aspects of the energy flux reconstruction are given in Jochum *et al*. (2021). I also calculated basic descriptors of food-web topology: connectance (proportion of realised links), graph centralisation (concentration of interactions around central nodes of the network; <u>Freeman, 1978)</u> and modularity (presence of interaction clusters; Fig. 8C) (Newman & Girvan, 2004; Laigle *et al*., 2018).

**Figure 8.**
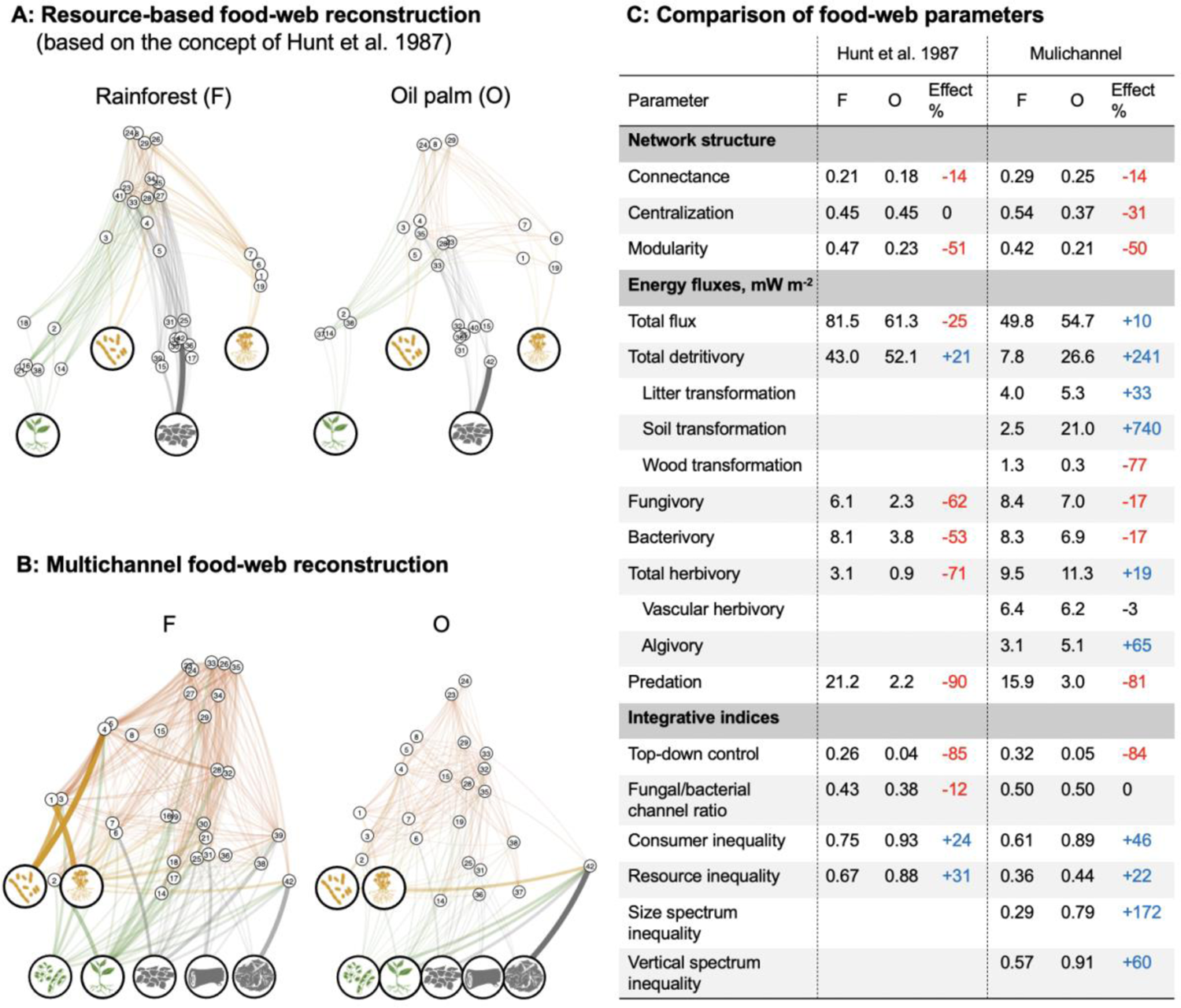
Comparing network topologies and energy flux-based indicators in traditional and multichannel soil food-web reconstructions. The comparison is based on the empirical data for nematodes, mesofauna and macrofauna collected from rainforests (F) and oil palm plantations (O) in Sumatra, Indonesia (Krashevska, 2019; Potapov *et al*., 2019b). Resource-based reconstruction is based on the ideas that primary consumers in belowground food webs diverge in their feeding preferences and cluster in resource-based energy channels, such as fungal, bacterial, root (plant) and detrital, that are coupled by predators (Hunt *et al*., 1987; Moore *et al*., 2005). Network nodes are ordered according to resource used (x axis) and trophic position (y axis) (**A**). Multichannel reconstruction captures widespread generalism in resource preferences with resource-based energy channels being to a large extent reticulated (Digel *et al*., 2014; Wolkovich, 2016) and consumers clustered also in body size and spatial energy channels (Potapov *et al*., 2021b); see above). Network nodes are ordered according to body mass (x axis) and trophic position (y axis) (**B**). Colours of the network edges highlight predation (dark orange), brown (grey and dark yellow) and green energy channels (green). (**C**) Comparing network topology, energy fluxes and integrative indicators of the two reconstructions. The difference in indicators between oil palm plantations and rainforests is shown as the percentage change (effect %); red denotes a reduction in plantations, blue denotes an increase in plantations, black denotes no considerable change (effect <10%). The two reconstructions resulted not only in different absolute estimations of functional indicators but also in different effects on them of oil palm cultivation.

The absolute estimators of virtually all calculated network parameters and functional indicators are different between the two reconstructions, highlighting the importance of network topology in food-web analysis (Fig. 8C). The multichannel reconstruction resulted in higher network connectance and a slightly lower network modularity, thus reflecting reticulated energy channels in the soil food-web (Fig. 8B). The multichannel reconstruction also resulted in a lower total energy flux estimation, higher estimations of herbivory and microbivory and a lower estimation of detritivory. These differences reflected trophic level omnivory of soil consumers – detritivores feed in large on microorganisms rather than the dead plant material itself (Swift *et al*., 1979; Larsen *et al*., 2016; Steffan *et al*., 2017; Potapov *et al*., 2019e). This increases assimilation efficiency and reduces the estimations of total energy flux and detritivory in the sense of dead plant material feeding (Fig. 2).

Effects of the land-use change from rainforest to plantations differed in magnitude and sometimes direction for several indicators calculated between the two reconstructions (Fig. 8C). Although the total energy flux decreased on plantations according to the resource-based reconstruction, it slightly increased according to the multichannel reconstruction. The total energy flux change in the multichannel reconstruction matched previous estimates (Potapov *et al*., 2019b). Bacterivory, fungivory and total herbivory were reduced by 53-71% in plantations according to the resource-based reconstruction, but the reduction was much smaller according to the multichannel reconstruction. There was an increase in herbivory of 19% (Fig. 8C). This difference reflects the fact that large detritivores (earthworms in plantations in this case) can feed on multiple resources and thus in part trophically compensate for the decline in the abundance of specialised bacterial, fungal and plant-feeding fauna. Again, the results of the multichannel reconstruction better reflected empirical data, showing that conversion of rainforest into oil palm plantations increases the plant energy channel in soil food webs (Klarner *et al*., 2017; Susanti *et al*., 2019) and reduces the bacterial, but not the fungal, energy channel (Susanti *et al*., 2019). In fact, resource-based reconstruction showed that the ratio of fungal to bacterial energy channelling is reduced in oil palm plantations. This contradicts empirical data from several invertebrate taxonomic groups (Susanti *et al*., 2019).

In both reconstructions, the estimates of detritivory were higher in oil palm plantations, despite the lower directly measured litter decomposition in this system compared to that in rainforest (Krashevska *et al*., 2018). However, in contrast to the resource-based reconstruction, it was possible with the multichannel reconstruction to distinguish divergent patterns in litter, soil and wood transformation processes (Fig. 8C). A strong decline in the energy flux related to wood transformation suggests that this index may reflect decomposition of recalcitrant organic matter and may be better related to overall litter decomposition (Table 2). A similar decline in both reconstructions was also observed for total predation. However, the traditional approach merged predators into a single pool, whereas the multichannel approach made it possible to distinguish between predators, coupling different resource-size- and spatial energy channels. Network graphs (Fig. 8B) illustrate a large reduction of large-sized predators in the soil food webs of oil palm plantations (x axis reflects body mass), clarifying the mechanisms of predation decline that was observed in previous reconstructions (Barnes *et al*., 2014; Potapov *et al*., 2019b).

Gini inequality indices further reflected unbalanced energy channelling along the size- and spatial food-web dimensions, thus reflecting not only general changes in food-web energetics but also the mechanisms behind these changes (e.g. resource shift or changes in ecosystem structure). This was possible only using the multichannel reconstruction. In the example given, multichannel reconstruction made it possible to reveal the strong increase in inequality across the size spectrum, vertical spectrum and among consumers (i.e. the dominance of the earthworm node that have a large body mass and inhabit soil in plantations; Fig. 2C). Unchanged total energy flux and increased energetic inequality suggest that food-web multifunctionality and stability is compromised in plantation systems (Table 2).

## IV. Evaluation and the way forward

### (1) Critical evaluation and knowledge gaps

My main goal was to make soil food-web reconstructions more realistic and make them more accessible by providing a reproducible analytical framework and by linking energy channelling to various consumer-driven ecosystem functions. I connected soil protists, invertebrates and vertebrates into a single interaction network, allowing for the simultaneous quantitative comparison of their ecosystem roles. I claim the reconstruction to be scalable across different food-web compartments and different spatial scales. I also believe that the approach can deliver informative ecosystem indicators, some of which could be used to include consumers in biogeochemical models. Nevertheless, these claims all have some issues that must be considered.

Of these issues, the most important is that food-web reconstruction is only as good as our knowledge of soil animal biology. And this knowledge is still fragmentary (Geisen *et al*., 2019). The feeding habits of many groups have not been validated with rigorous empirical approaches and most of the information comes from well-studied species in well-studied ecosystems (Potapov, Tiunov, & Scheu, 2019d; Velazco *et al*., 2021). Shifts in feeding habits is one of the response mechanisms of some soil consumers to environmental changes (Krause *et al*., 2019), which is hard to include in the food-web reconstruction without direct assessment of trophic interactions. Food webs in different ecosystems are assembled from different species and trophic guilds but the same trophic guild may also shift its ecosystem role if major shifts occur in the environment (Susanti *et al*., 2019). These systematic between-ecosystem variations that are able to bias comparison based on feeding guilds should be further explored. Another critical aspect of the food-web reconstruction is defining preferences for omnivorous species that feed both on basal resources and other consumers (Jochum *et al*., 2021). Biomass-defined preferences overestimate the contribution of basal resources to the diet due to their omnipresence. In my reconstruction I have manually adjusted feeding on basal/animal resources according to existing knowledge (Potapov *et al*., [bioRxiv]). This is more realistic than the common practice of assigning equal preferences to all resources (Barnes *et al*., 2014; Jochum *et al*., 2021). Nevertheless, such decisions are able to propagate into biased energy flux estimations (Jochum *et al*., 2021) and call for further validation of the feeding preferences across different trophic guilds. Finally, several other assumptions behind the reconstruction, such as protective mechanisms and PPMRs, should be tested and the effect of different traits should be quantitatively assessed (Peschel *et al*., 2006; Schneider & Maraun, 2009; Eitzinger *et al*., 2018). Despite all these uncertainties, I achieved a realistic reconstruction based on relatively simple rules without apparent contradictions with what is already commonly known. Furthermore, this reconstruction is open to further improvement. Importantly, assumptions about food-web topology in traditional food-web reconstructions (Hunt *et al*., 1987) have never been critically tested and often are not in line with empirical data (Digel *et al*., 2014; Geisen, 2016; Wolkovich, 2016).

The multichannel reconstruction is scalable and can be applied across or within food-web compartments. However, the approach has less power if only few species from one compartment or size class are considered because of the potentially large effect of species-specific interactions that might be overlooked. This uncertainty is in part counteracted in reconstructions across food-web compartments by the wide range of trophic guilds and taxa considered. Body mass range is particularly important for the reconstruction because of allometric predator-prey interactions. Unfortunately, many studies target only a small component of soil food-web. I want to change this fragmentary vision to a more holistic approach and here I provided a tool for describing and quantifying entire soil communities, from microbes to vertebrates. Despite being demanding of labour and requiring a diverse toolbox, complex assessment of animal communities provides unique opportunities for understanding ecosystem functioning and is feasible with collaborative research. Such assessments will become more accessible with the development of e.g. image analysis tools that provide taxonomic identification together with body size and biomass estimations (Ärje *et al*., 2020). Revealing mechanisms that control energy channelling in soil food webs and testing how different energetic configurations of soil food webs affect ecosystem processes in controlled experiments can deliver holistic food-web indicators. An empirical validation of the links between food-web structure and ecosystem functioning using the multichannel reconstruction is the next crucial step for including holistic indicators of consumer communities in biogeochemical models.

### (2) Expanding dimensionality

While presenting the multichannel reconstruction, I focused on the dimensionality of soil food webs across resources, size classes and soil layers. Future developments may introduce more dimensions, such as the temporal dimension. Although inhabiting the same layer and having similar size, some species or functional guilds may have limited interaction due to differentiation in their daily or seasonal activity patterns. For example, most amphibians are active at night whereas reptiles are often active in the daytime. Many holometabolous insect groups are active in soils seasonally, before their aboveground-living imago emerge.

Furthermore, in the spatial dimension I have focused on the vertical stratification of soil food webs but it would also be possible to consider the horizontal distribution. Soil food webs are clustered around microsites with high activity, i.e. ‘hotspots’, such as drilosphere and rhizosphere (Thakur *et al*., 2020) and local food webs are connected through mobile surface- and aboveground-dwelling consumers into meta-webs (Mougi & Kondoh, 2016; Hirt *et al*., 2018), which also can be quantified with energy fluxes.

Among specific traits, elemental composition can be considered in the food selection (Buchkowski & Lindo, 2020). Node-specific cannibalistic interactions could be quantified and incorporated. Trait matching algorithms, such as visual hunting versus camouflage protection, could be accounted for. Individual trophic flexibility varies among species (Krause *et al*., 2019) and defines food choice under different settings, e.g. depending on resource availability. This characteristic can be included in the model to assign node-specific trophic flexibility. In fact, any functional trait can be incorporated in the multichannel reconstruction to improve the prediction of trophic interactions or summarise certain trophic functions.

Increase in complexity of the reconstruction due to the incorporation of many traits will not necessarily lead to a proportional increase in the complexity of the calculations if data and programming code will be openly shared (Table S1 and Appendix S3).

### (3) Beyond the soil

Throughout the text I focused on soil food webs. I did so because these food webs are cryptic and less understood that those in water or above ground and because I am able to validate the approach, in part, with my own knowledge. Nevertheless, the energy flux approach can be applied across ecosystem types (Barnes *et al*., 2018) and the multichannel reconstruction can be flexibly expanded to include aboveground, freshwater and marine consumers. By introducing the spatial aspect of habitat preference in the network reconstruction, I opened the opportunity of quantifying energy exchange between ecosystem compartments based on energy fluxes through the nodes that belong to different ecosystem compartments or moving between them. Network stability and motifs, total fluxes, channel structures, trophic hierarchies and related ecosystem functions can be statistically compared among food-web compartments, ecosystem types and ecosystems.

## V. Conclusions

1. Soil food webs are organized along several dimensions, according to the resource use, body size classes and environmental heterogeneity in space. Until now, soil food-web reconstructions did not consider these dimensions together and a reproducible approach to describe widespread omnivory and multichannel feeding of soil-associated consumers has not been suggested.
2. In the present paper, I describe the multichannel reconstruction of soil food webs based on generic food-web organization principles and functional classification of consumers, including protists, invertebrates and vertebrates. The reconstruction can be applied using data on trophic guild abundances, even if the trophic links were not measured directly in the field.
3. Using the energy flux approach and the multichannel reconstruction, I further summarised existing and novel quantitative indicators of trophic functions and food-web stability. Suggested indicators can be used to assess trophic multifunctionality (analogous to ecosystem multifunctionality) and a wide spectrum of single trophic and ecosytem functions, including herbivory, detritus transformation and translocation, microbial grazing and dispersal and top-down control.
4. The multichannel reconstruction differs from traditional food-web reconstruction in estimated network topology parameters and calculated trophic functions. The multichannel reconstruction better matched some of the independently measured ecosystem functions and food-web parameters, but this conclusion is tentative, since systematic research is needed to validate the approach. An advantage of the multichannel reconstruction is that it can describe multiple aspects of food-web functioning, providing higher resolution and more mechanistic understanding of observed food-web variations than traditional food-web reconstructions.
5. Further validation, development and application of the multichannel reconstruction will allow us to achieve realistic and holistic functional description of soil consumer communities in different ecosystems. More characteristics of trophic guilds can be flexibly incorporated in the multichannel approach, which bridges this way food-web ecology with functional trait ecology. Suggested functional indicators could be further used to depict the contribution of animals and protists in the processes of organic matter transformation and nutrient mineralisation and include into biogeochemical models the top-down control of ecosystem functioning by consumers in soil on the local, regional and even the global scale.

## Acknowledgements

The work was supported by the Deutsche Forschungsgemeinschaft (DFG, German Research Foundation) project number 192626868 – SFB 990 in the framework of the collaborative German - Indonesian research project CRC990 - EFForTS. Initial idea of the review was developed in the framework of the mobility program 57448388, co-funded by Czech Academy of Sciences (CAS) and Deutscher Akademischer Austauschdienst (DAAD). Work was also in part supported by the Alexander von Humboldt foundation in the framework of the Research group linkage programme “Structure and functioning of belowground food webs across temperate and tropical ecosystems”. I am grateful to Malte Jochum for all the discussions about the energy flux approach and his support during the preparation of this manuscript. I am also grateful to Alexei Tiunov and Stefan Scheu for introducing me into the cryptic world of belowground food webs. Special gratitude goes to Svenja Meyer for preparation of the animal silhouettes. The manuscript was edited during preparation by Dr AJ Davis (English Experience Language Services, Göttingen, Germany).

## Supplementary materials

Appendix S1: Table S1 list of trophic guilds used in the exemplar reconstruction Appendix S2 – Textual description of food-web reconstruction

Appendix S2: Table S1 Correction coefficients for predator traits Appendix S2: Table S2 – Correction coefficients for prey traits Appendix S3 – R code for multichannel food-web reconstruction Supplementary Table S1 – descriptions of the data in the Table S2 Supplementary Table S2 – the full list of trophic guilds with traits

### Appendix S1: Guild table

**Appendix S1: Table S1.**
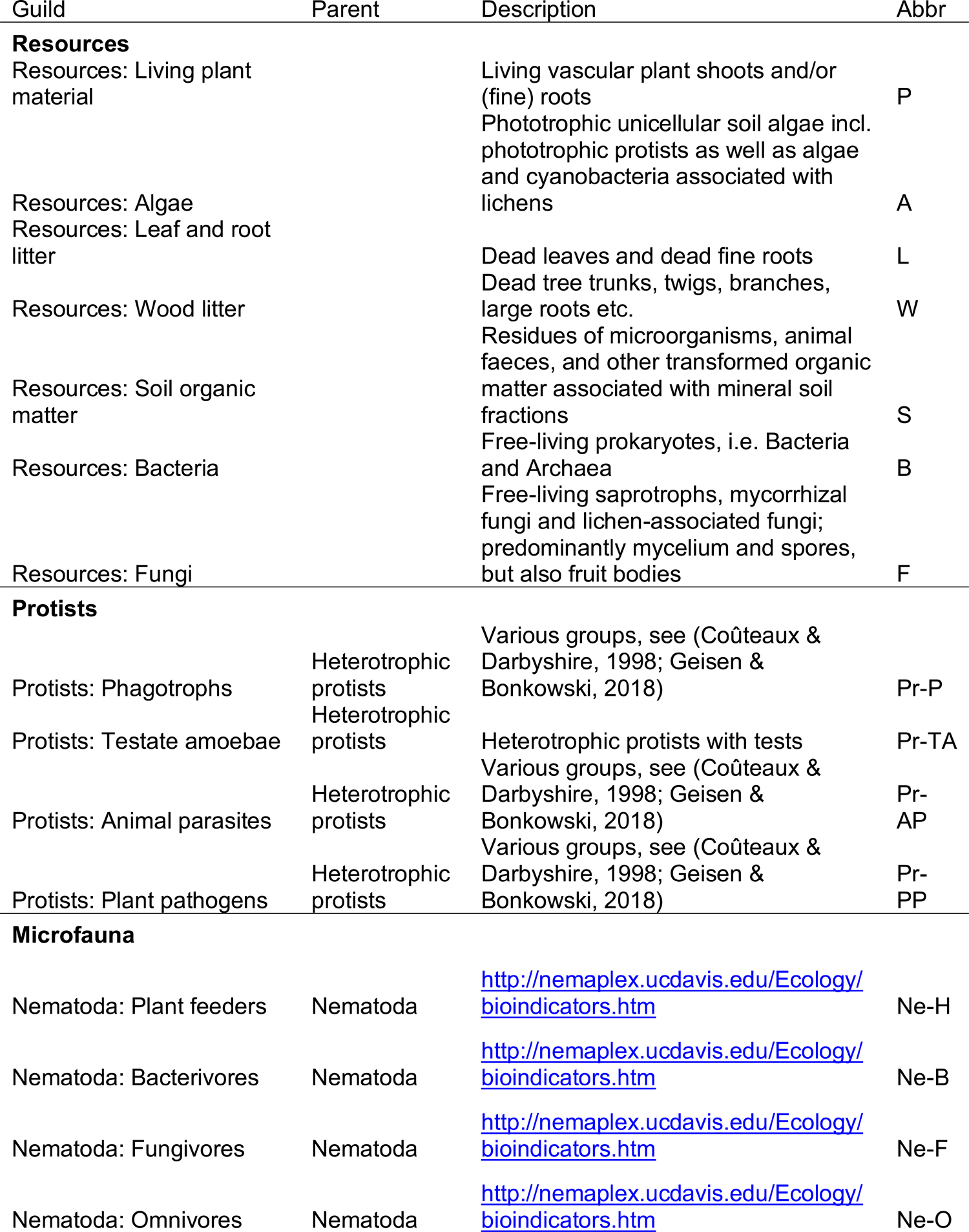
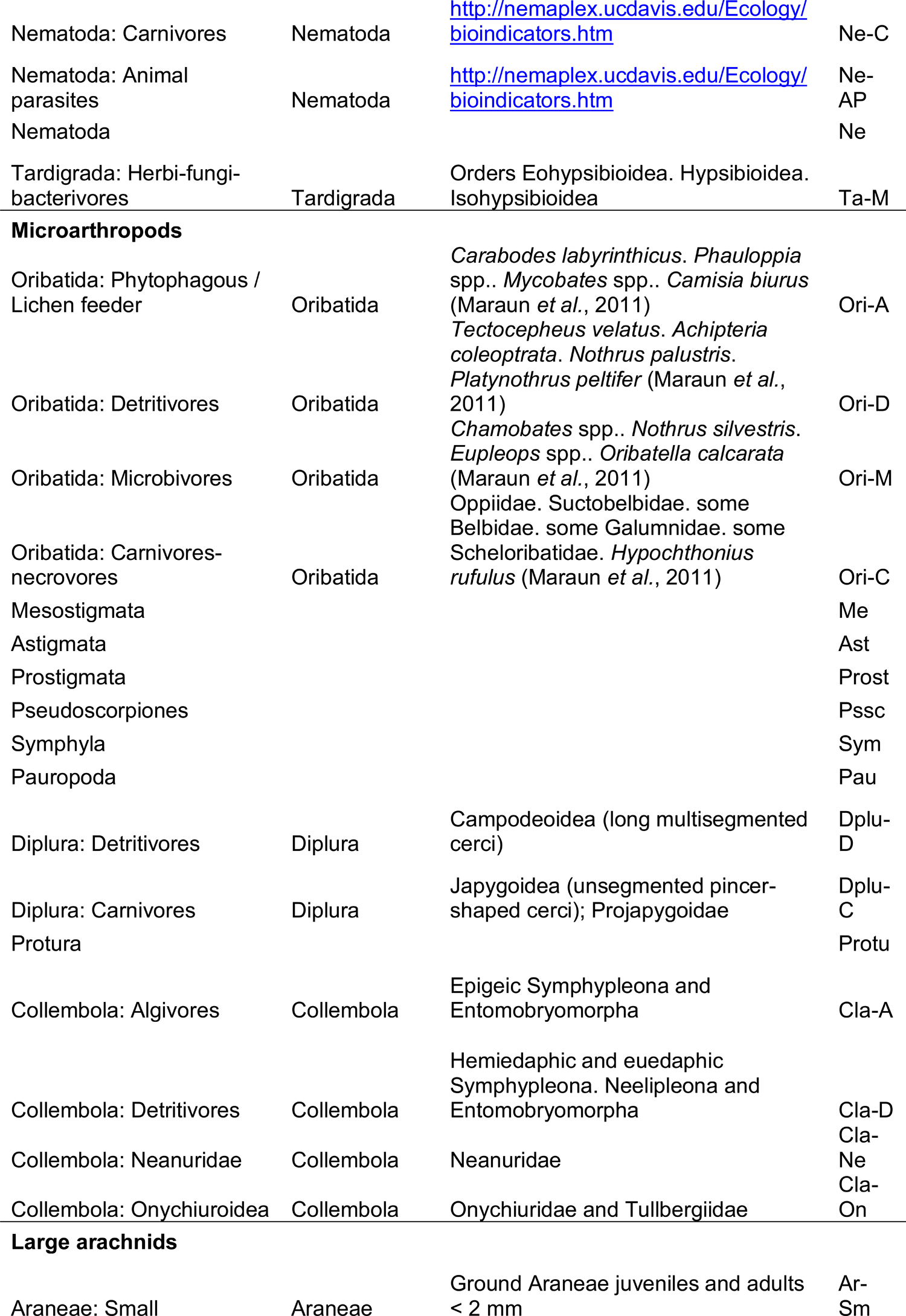
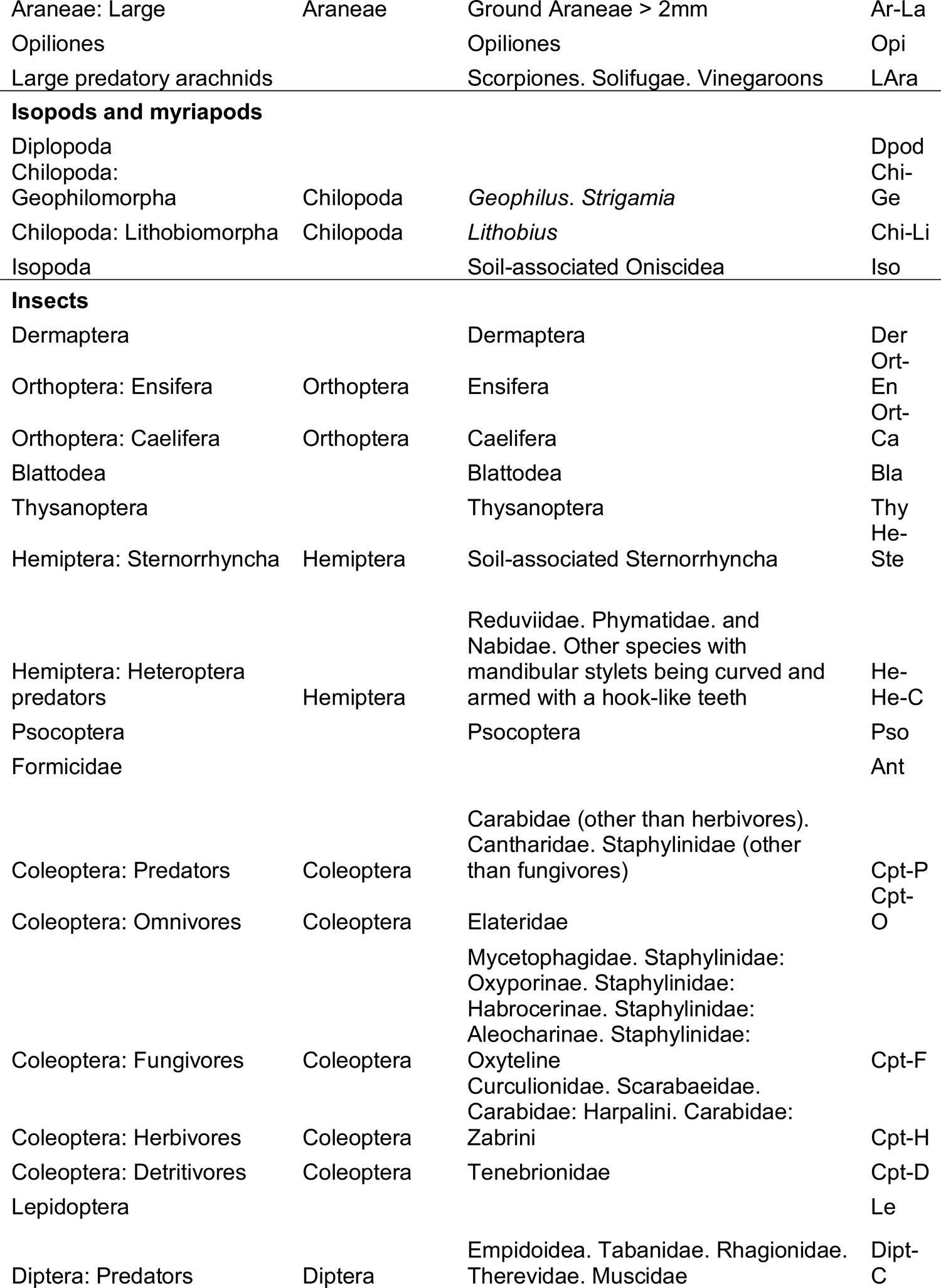
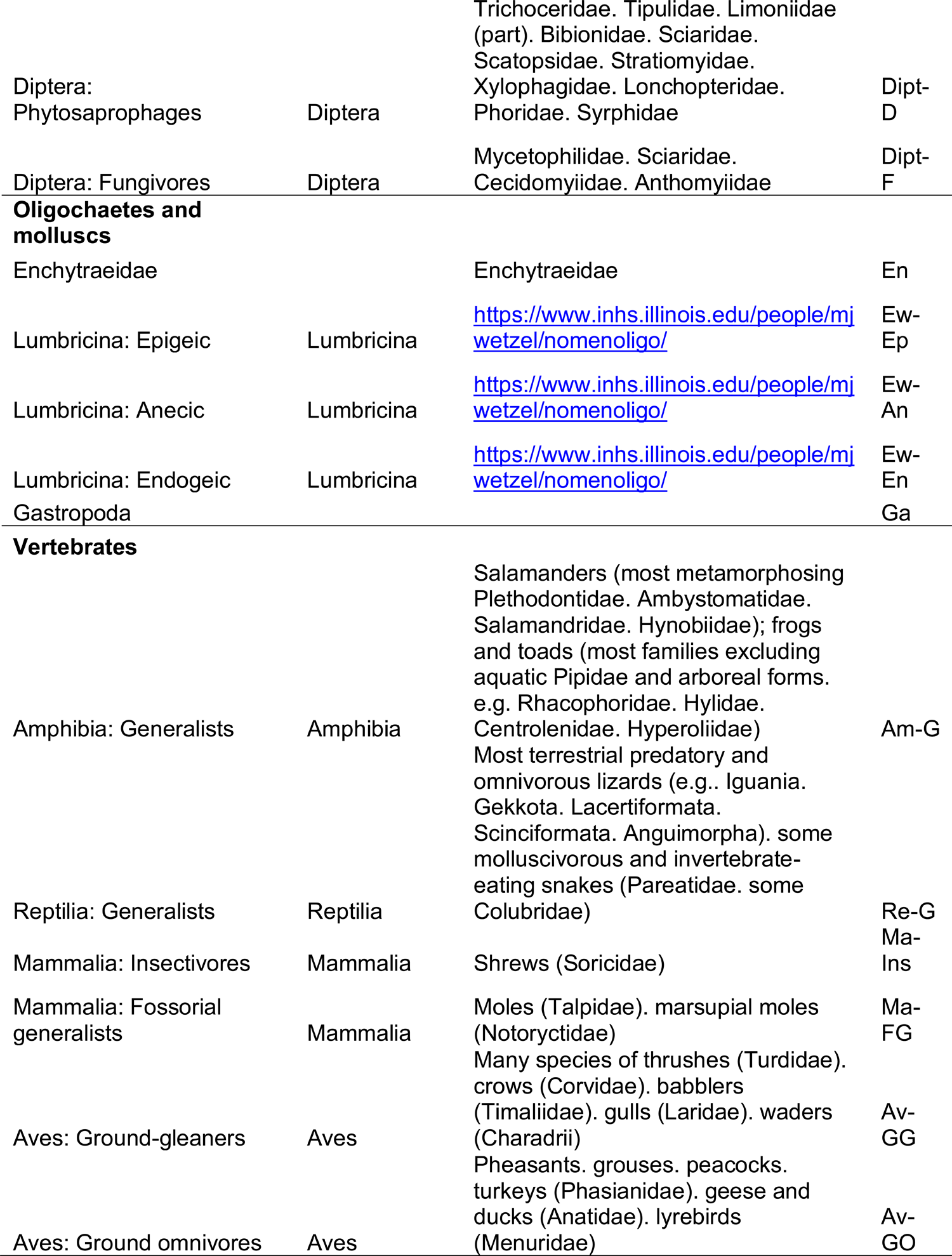
List of trophic guilds used in the exemplar reconstruction. In all cases, taxonomic groups refer to the soil-associated species only. Parent – guild of a higher hierarchical level. Abbr – abbreviation.

### Appendix S2: Food-web reconstruction

Food-web reconstruction was based on the trophic guild classification and other data provided in the Table S1; 7 resource and 66 consumer nodes marked in the “Example” column were used. The following parameters were used for each node: (1) feeding preferences to basal resources and predation capability; (2) mean and standard deviation of the log_10_-transformed living body mass; (3) coefficients for optimum PPMR and PPMR width based on predator traits; (4) protection coefficient based on prey protective traits; (5) occurrence in soil, litter, surface and aboveground (vertical distribution). Feeding preferences and vertical distribution were coded as 0 (nearly absent), 0.5 (occasionally/supplementary) and 1 (present). The following default assumptions were used for the exemplar reconstruction: optimum PPMR was set to 100 (PPMRopt = log10(100)) (Brose *et al*., 2008), body mass range of the optimum prey was set to be the same as the body mass range of the predator (PPMRwidth = 1), feeding on the additional resources (coded as 0.5) was assumed to be 20% of that on main resources (coded as 1). In case of omnivory, i.e. developed predation capabilities and feeding on any of basal resources (both coded as 1), animal diet was assumed to represent 50% of the node budget (Barnes *et al*., 2014). To reconstruct the food-web, I generated a set of adjacency matrices based on different parameters with food objects in rows and consumers in columns and multiplied them.

First, I generated the resource adjacency matrix. For each node, feeding preferences were distributed across all possible basal resources and other consumers assuming that additional resources are five times less important than the main resources (see above).

Omnivores were assumed to feed 50% on animal food and 50% on all other basal resources. All preferences for each node (columns) summed up to 1.

Second, I generated the allometric adjacency matrix. For each node, I modelled body size distribution using the mean and standard deviation of living body mass and assuming it follows log-normal pattern, which generally observed both across species (Hutchinson & MacArthur, 1959; Blackburn & Gaston, 1994; Allen *et al*., 2006) and within species (Stead *et al*., 2005; Potapov *et al*., 2021b). Then I calculated overlaps of the size distributions, correcting PPMR for all potential predator-prey interactions (Fig. 4). Overlap proportion [0;1] was used as the proxy of interaction strength. I corrected optimum PPMR for corresponding trait-based coefficients.

Third, I generated the protection matrix. I filled the matrix with ones and multiplied it or the protection coefficient for each node by rows. Coefficients varied from 0.112 to 1.

Fourth, I generated the spatial adjacency matrix. I calculated pairwise Bray-Curtis dissimilarities for all nodes based on the vertical distribution in soil, litter, surface and aboveground. Reverse dissimilarity [0;1] as the measure of spatial overlap was used as the proxy of interaction strength.

The resource, allometric, protection and spatial matrices were multiplied to generate the final adjacency matrix that represented strengths of feeding interactions among food-web nodes. The matrix together with metabolic losses and biomasses was used in the *fluxweb* package to calculate energy fluxes (Gauzens *et al*., 2019). Due to a lack of an community with available biomasses from protists to vertebrates from which I could draw example biomasses, I ignored biomasses and metabolic losses in the first food-web reconstruction and used initial interaction strengths as the proxy of energy fluxes. Thus, the results reported for the first food-web reconstruction are unrealistic and all numbers are shown only to illustrate the concept.

**Appendix S2: Table S1.**
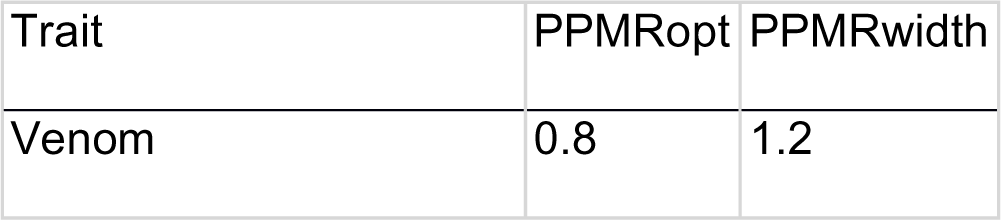
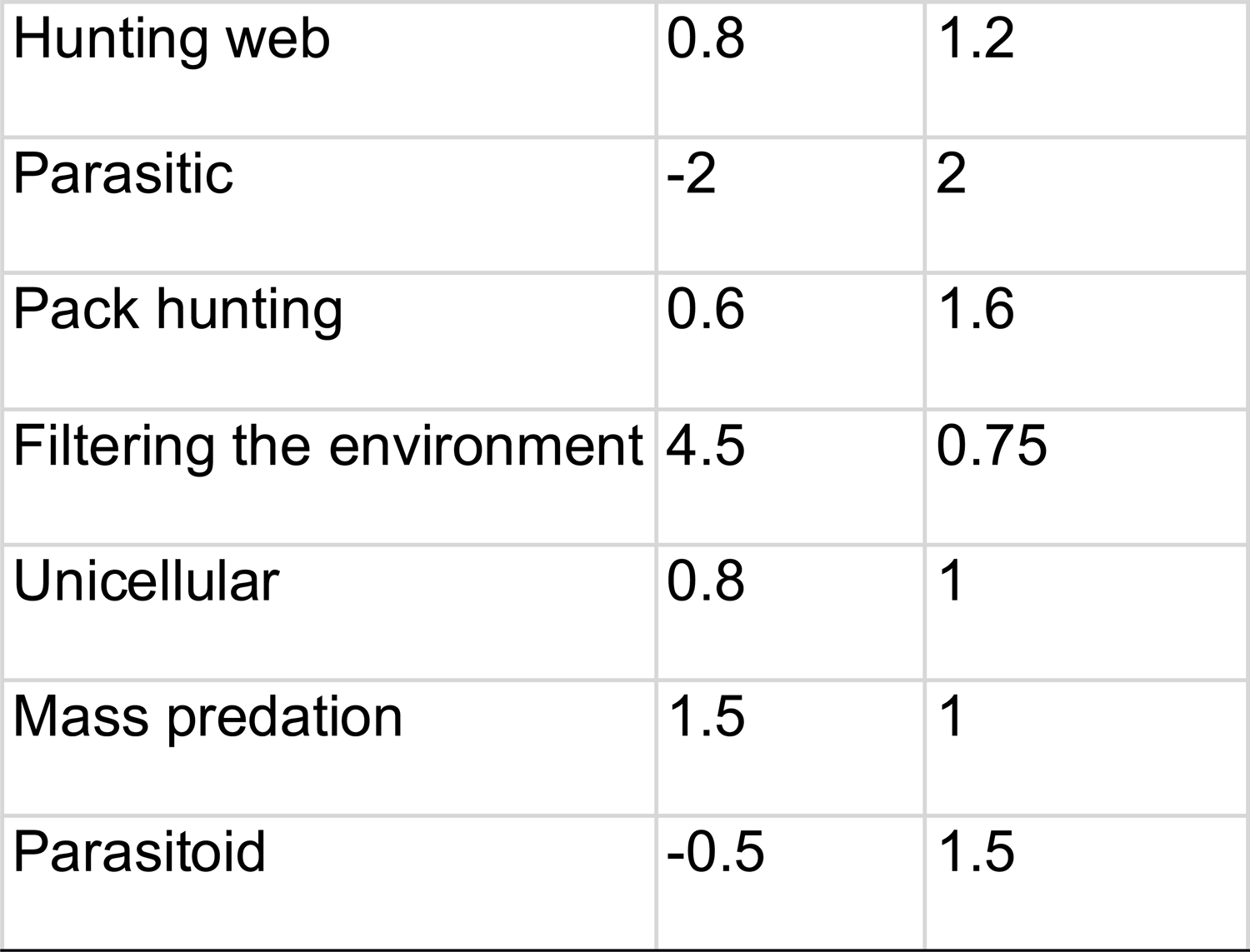
**Correction coefficients for predator traits used in the food-web reconstruction**. Values are given as multiplication coefficients and are based on my expert opinion mostly to illustrate the concept. Each trait may reduce or increase optimum predator-prey mass ratio (PPMRopt) and also reduce or increase the width of the prey mass range (PPMRwidth).

| Trait | PPMR<sub>opt</sub> | PPMR<sub>width</sub> |
| --- | --- | --- |
| Venom | 0.8 | 1.2 |
| Hunting web | 0.8 | 1.2 |
| Parasitic | -2 | 2 |
| Pack hunting | 0.6 | 1.6 |
| Filtering the environment | 4.5 | 0.75 |
| Unicellular | 0.8 | 1 |
| Mass predation | 1.5 | 1 |
| Parasitoid | -0.5 | 1.5 |

**Appendix S2: Table S2.** **Correction coefficients for prey traits used in the food-web reconstruction**. Values are given as multiplication coefficients and are based on my expert opinion mostly for the concept illustration. Each trait protects prey and thus decreases predation rate (the smaller the coefficient, the bigger the effect).

| Trait | Protection |
| --- | --- |
| Carnivore | 0.7 |
| Agility (e.g. jumping ) | 0.7 |
| Moderate physical protection (e.g. strong cuticle) | 0.7 |
| Strong physical protection (e.g. shell) | 0.4 |
| Protective metabolites | 0.7 |
| Protective metabolites (strong) | 0.4 |
| Parasitic | 0.2 |

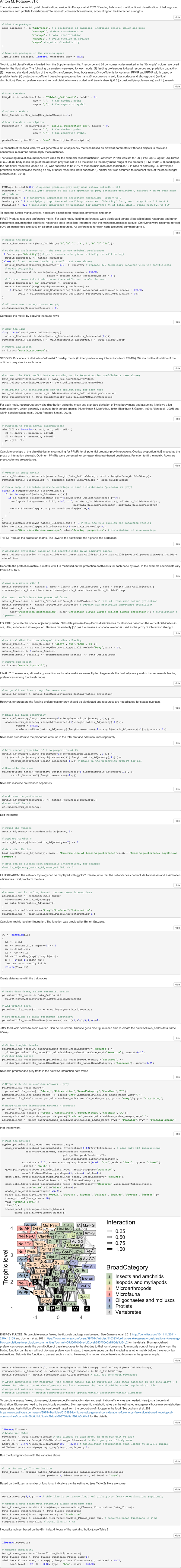
Multichannel food-web reconstruction.

## Notes

### Competing Interest Statement

The authors have declared no competing interest.

## References

1. Allen, C.R., Garmestani, A.S., Havlicek, T.D., Marquet, P.A., Peterson, G.D., Restrepo, C., Stow, C.A. & Weeks, B.E. (2006) Patterns in body mass distributions: sifting among alternative hypotheses. Ecology Letters 9, 630–643.

2. Allesina, S., Alonso, D. & Pascual, M. (2008) A General Model for Food Web Structure. Science 320, 658–661.

3. Anderson, J.M. (1975) The enigma of soil animal species diversity. In Progress in soil zoology pp. 51–58. Springer.

4. Ärje, J., Melvad, C., Jeppesen, M.R., Madsen, S.A., Raitoharju, J., Rasmussen, M.S., Iosifidis, A., Tirronen, V., Gabbouj, M., Meissner, K. & Høye, T.T. (2020) Automatic image-based identification and biomass estimation of invertebrates. Methods in Ecology and Evolution 11, 922–931.

5. Averill, C., Turner, B.L. & Finzi, A.C. (2014) Mycorrhiza-mediated competition between plants and decomposers drives soil carbon storage. Nature 505, 543–545.

6. Barnes, A.D., Jochum, M., Lefcheck, J.S., Eisenhauer, N., Scherber, C., O’connor, M.I., De Ruiter, P. & Brose, U. (2018) Energy flux: The link between multitrophic biodiversity and ecosystem functioning. Trends in Ecology & Evolution 33, 186–197.

7. Barnes, A.D., Jochum, M., Mumme, S., Haneda, N.F., Farajallah, A., Widarto, T.H. & Brose, U. (2014) Consequences of tropical land use for multitrophic biodiversity and ecosystem functioning. Nature Communications 5, 5351.

8. Barnes, A.D., Scherber, C., Brose, U., Borer, E.T., Ebeling, A., Gauzens, B., Giling, D.P., Hines, J., Isbell, F., Ristok, C., Tilman, D., Weisser, W.W. & Eisenhauer, N. (2020) Biodiversity enhances the multitrophic control of arthropod herbivory. Science Advances 6. American Association for the Advancement of Science.

9. Bauer, T. & Pfeiffer, M. (1991) ‘Shooting’springtails with a sticky rod: the flexible hunting behaviour of Stenus comma (Coleoptera; Staphylinidae) and the counter-strategies of its prey. Animal Behaviour 41, 819–828. Elsevier.

10. Berg, M.P. & Bengtsson, J. (2007) Temporal and spatial variability in soil food web structure. Oikos 116, 1789–1804.

11. Blackburn, T.M. & Gaston, K.J. (1994) Animal body size distributions: patterns, mechanisms and implications. Trends in Ecology & Evolution 9, 471–474. Elsevier.

12. Blanchard, J.L., Heneghan, R.F., Everett, J.D., Trebilco, R. & Richardson, A.J. (2017) From bacteria to whales: using functional size spectra to model marine ecosystems. Trends in ecology & evolution 32, 174–186.

13. Bluhm, S.L., Potapov, A.M., Shrubovych, J., Ammerschubert, S., Polle, A. & Scheu, S. (2019) Protura are unique: first evidence of specialized feeding on ectomycorrhizal fungi in soil invertebrates. BMC Ecology 19.

14. Bouché, M.B. (1977) Strategies lombriciennes. Ecological Bulletins, 122–132.

15. Bradford, M.A. (2016) Re-visioning soil food webs. Soil Biology and Biochemistry 102, 1–3.

16. Bradford, M.A., Jones, T.H., Bardgett, R.D., Black, H.I., Boag, B., Bonkowski, M., Cook, R., Eggers, T., Gange, A.C., Grayston, S.J., & Others (2002) Impacts of soil faunal community composition on model grassland ecosystems. Science 298, 615–618.

17. Bradford, M.A., Warren Ii, R.J., Baldrian, P., Crowther, T.W., Maynard, D.S., Oldfield, E.E., Wieder, W.R., Wood, S.A. & King, J.R. (2014) Climate fails to predict wood decomposition at regional scales. Nature Climate Change 4, 625–630.

18. Briones, M.J.I. (2014) Soil fauna and soil functions: a jigsaw puzzle. Frontiers in Environmental Science 2.

19. Brose, U., Archambault, P., Barnes, A.D., Bersier, L.-F., Boy, T., Canning-Clode, J., Conti, E., Dias, M., Digel, C., Dissanayake, A., Flores, A.A.V., Fussmann, K., Gauzens, B., Gray, C., Häussler, J., et al. (2019) Predator traits determine food-web architecture across ecosystems. Nature Ecology & Evolution 3, 919–927.

20. Brose, U., Ehnes, R.B., Rall, B.C., Vucic-Pestic, O., Berlow, E.L. & Scheu, S. (2008) Foraging theory predicts predator-prey energy fluxes. Journal of Animal Ecology 77, 1072–1078.

21. Brose, U. & Scheu, S. (2014) Into darkness: unravelling the structure of soil food webs. Oikos 123, 1153–1156.

22. Brousseau, P.-M., Gravel, D. & Handa, I.T. (2018) Trait matching and phylogeny as predictors of predator-prey interactions involving ground beetles. Functional Ecology 32, 192–202.

23. Brown, J.H., Gillooly, J.F., Allen, A.P., Savage, V.M. & West, G.B. (2004) Toward a metabolic theory of ecology. Ecology 85, 1771–1789.

24. Brussaard, L. (1998) Soil fauna, guilds, functional groups and ecosystem processes. Applied Soil Ecology 9, 123–135.

25. Buchkowski, R.W. & Lindo, Z. (2020) Stoichiometric and structural uncertainty in soil food web models. Functional Ecology n/a.

26. Cardoso, P., Pekár, S., Jocqué, R. & Coddington, J.A. (2011) Global Patterns of Guild Composition and Functional Diversity of Spiders. PLoS ONE 6, e21710.

27. Cebrian, J. (1999) Patterns in the fate of production in plant communities. The American Naturalist 154, 449–468.

28. Cerdá, X. & Dejean, A. (2011) Predation by ants on arthropods and other animals. In Predation in the Hymenoptera: An evolutionary perspective (ed C. POLIDORI), pp. 39–78.

29. Ceriani, L. & Verme, P. (2012) The origins of the Gini index: extracts from Variabilità e Mutabilità (1912) by Corrado Gini. The Journal of Economic Inequality 10, 421–443. Springer.

30. Chertov, O., Komarov, A., Shaw, C., Bykhovets, S., Frolov, P., Shanin, V., Grabarnik, P., Priputina, I., Zubkova, E. & Shashkov, M. (2017) Romul_Hum—A model of soil organic matter formation coupling with soil biota activity. II. Parameterisation of the soil food web biota activity. Ecological Modelling 345, 125–139.

31. Clemmensen, K.E., Bahr, A., Ovaskainen, O., Dahlberg, A., Ekblad, A., Wallander, H., Stenlid, J., Finlay, R.D., Wardle, D.A. & Lindahl, B.D. (2013) Roots and Associated Fungi Drive Long-Term Carbon Sequestration in Boreal Forest. Science 339, 1615–1618.

32. Cohen, J.E., Pimm, S.L., Yodzis, P. & Saldaña, J. (1993) Body sizes of animal predators and animal prey in food webs. Journal of Animal Ecology 62, 67–78.

33. Coleman, D.C., Callaham, M.A. & Crossley Jr, D. (2017) Fundamentals of soil ecology, 3d edition. Academic press.

34. Coûteaux, M.-M. & Darbyshire, J.F. (1998) Functional diversity amongst soil protozoa. Applied Soil Ecology 10, 229–237.

35. Crowther, T.W., Van Den Hoogen, J., Wan, J., Mayes, M.A., Keiser, A.D., Mo, L., Averill, C. & Maynard, D.S. (2019) The global soil community and its influence on biogeochemistry. Science 365.

36. Deckmyn, G., Flores, O., Mayer, M., Domene, X., Schnepf, A., Kuka, K., Looy, K.V., Rasse, D.P., Briones, M.J.I., Barot, S., Berg, M., Vanguelova, E., Ostonen, I., Vereecken, H., Suz, L.M., et al. (2020) KEYLINK: towards a more integrative soil representation for inclusion in ecosystem scale models. I. review and model concept. PeerJ 8, e9750. PeerJ Inc.

37. Delgado-Baquerizo, M., Reich, P.B., Trivedi, C., Eldridge, D.J., Abades, S., Alfaro, F.D., Bastida, F., Berhe, A.A., Cutler, N.A., Gallardo, A., García-Velázquez, L., Hart, S.C., Hayes, P.E., He, J.-Z., Hseu, Z.-Y., et al. (2020) Multiple elements of soil biodiversity drive ecosystem functions across biomes. Nature Ecology & Evolution 4, 210–220.

38. Digel, C., Curtsdotter, A., Riede, J., Klarner, B. & Brose, U. (2014) Unravelling the complex structure of forest soil food webs: higher omnivory and more trophic levels. Oikos 123, 1157–1172.

39. Dowdy, W.W. (1944) The influence of temperature on vertical migration of invertebrates inhabiting different soil types. Ecology 25, 449–460.

40. Ehnes, R.B., Pollierer, M.M., Erdmann, G., Klarner, B., Eitzinger, B., Digel, C., Ott, D., Maraun, M., Scheu, S. & Brose, U. (2014) Lack of energetic equivalence in forest soil invertebrates. Ecology 95, 527–537.

41. Ehnes, R.B., Rall, B.C. & Brose, U. (2011) Phylogenetic grouping, curvature and metabolic scaling in terrestrial invertebrates: Invertebrate metabolism. Ecology Letters 14, 993– 1000.

42. Eisenbeis, G. & Wichard, W. (1987) Atlas on the biology of soil arthropodsEnglish version of original edition. Springer-Verlag, Berlin Heidelberg New York London Paris Tokyo.

43. Eitzinger, B., Rall, B.C., Traugott, M. & Scheu, S. (2018) Testing the validity of functional response models using molecular gut content analysis for prey choice in soil predators. Oikos.

44. Ellers, J., Berg, M.P., Dias, A.T.C., Fontana, S., Ooms, A. & Moretti, M. (2018) Diversity in form and function: Vertical distribution of soil fauna mediates multidimensional trait variation. Journal of Animal Ecology 87, 933–944.

45. Erktan, A., Or, D. & Scheu, S. (2020) The physical structure of soil: determinant and consequence of trophic interactions. Soil Biology & Biochemistry.

46. Ettema, C.H. & Wardle, D.A. (2002) Spatial soil ecology. Trends in ecology & evolution 17, 177–183.

47. Filser, J., Faber, J.H., Tiunov, A.V., Brussaard, L., Frouz, J., De Deyn, G., Uvarov, A.V., Berg, M.P., Lavelle, P., Loreau, M., Wall, D.H., Querner, P., Eijsackers, H. & Jiménez, J.J. (2016) Soil fauna: key to new carbon models. Soil 2, 565–582.

48. Freeman, L.C. (1978) Centrality in social networks conceptual clarification. Social networks 1, 215–239. North-Holland.

49. Fujii, S., Berg, M.P. & Cornelissen, J.H.C. (2020) Living Litter: Dynamic Trait Spectra Predict Fauna Composition. *Trends in Ecology & Evolution*, S0169534720301385.

50. Gauzens, B., Barnes, A., Giling, D.P., Hines, J., Jochum, M., Lefcheck, J.S., Rosenbaum, B., Wang, S. & Brose, U. (2019) *fluxweb* : An R package to easily estimate energy fluxes in food webs. Methods in Ecology and Evolution 10, 270–279.

51. Geisen, S. (2016) The bacterial-fungal energy channel concept challenged by enormous functional versatility of soil protists. Soil Biology and Biochemistry 102, 22–25.

52. Geisen, S. & Bonkowski, M. (2018) Methodological advances to study the diversity of soil protists and their functioning in soil food webs. Applied soil ecology 123, 328–333.

53. Geisen, S., Briones Mariaj. I., Gan, H., Behan-Pelletier, V.M., Friman, V.-P., Arjen De Groot, G., Hannula, S.E., Lindo, Z., Philippot, L., Tiunov, A.V. & Wall, D.H. (2019) A methodological framework to embrace soil biodiversity. Soil Biology and Biochemistry 136, 107536.

54. Ghilarov, M.S. (1949) Features of soil as a habitat and its importance in the evolution of insects. Russian Academy of Sciences, Moscow.

55. Gisin, H. (1943) Ökologie und Lebensgemeinschaften der Collembolen im schweizerischen Exkursionsgebiet Basels. A. Kundig, Geneve.

56. Gongalsky, K.B. (2021) Soil macrofauna: Study problems and perspectives. Soil Biology and Biochemistry 159, 108281.

57. Gongalsky, K.B., Zaitsev, A.S., Korobushkin, D.I., Saifutdinov, R.A., Butenko, K.O., Vries, F.T., Ekschmitt, K., Degtyarev, M.I., Gorbunova, A.Yu., Kostina, N.V., Rakhleeva, A.A., Shakhab, S.V., Yazrikova, T.E., Wolters, V. & Bardgett, R.D. (2021) Forest fire induces short-term shifts in soil food webs with consequences for carbon cycling. Ecology Letters 24, 438–450.

58. Grandy, A.S., Wieder, W.R., Wickings, K. & Kyker-Snowman, E. (2016) Beyond microbes: Are fauna the next frontier in soil biogeochemical models? Soil Biology and Biochemistry 102, 40–44.

59. Grass, I., Kubitza, C., Krishna, V.V., Corre, M.D., Mußhoff, O., Pütz, P., Drescher, J., Rembold, K., Ariyanti, E.S., Barnes, A.D., Brinkmann, N., Brose, U., Brümmer, B., Buchori, D., Daniel, R., et al. (2020) Trade-offs between multifunctionality and profit in tropical smallholder landscapes. Nature Communications 11, 1186.

60. Guerra, C.A., Heintz-Buschart, A., Sikorski, J., Chatzinotas, A., Guerrero-Ramírez, N., Cesarz, S., Beaumelle, L., Rillig, M.C., Maestre, F.T., Delgado-Baquerizo, M., Buscot, F., Overmann, J., Patoine, G., Phillips, H.R.P., Winter, M., et al. (2020) Blind spots in global soil biodiversity and ecosystem function research. Nature Communications 11, 3870.

61. Herberstein, M.E. (2011) Spider Behaviour: Flexibility and Versatility. Cambridge University Press.

62. Hines, J., Giling, D.P., Rzanny, M., Voigt, W., Meyer, S.T., Weisser, W.W., Eisenhauer, N. & Ebeling, A. (2019) A meta-food web for invertebrate species collected in a european grassland. *Ecology*, e02679.

63. Hines, J., Van Der Putten, W.H., De Deyn, G.B., Wagg, C., Voigt, W., Mulder, C., Weisser, W.W., Engel, J., Melian, C., Scheu, S., Birkhofer, K., Ebeling, A., Scherber, C. & Eisenhauer, N. (2015) Towards an Integration of Biodiversity– Ecosystem Functioning and Food Web Theory to Evaluate Relationships between Multiple Ecosystem Services. In Advances in Ecological Research pp. 161–199. Elsevier.

64. Hirt, M.R., Grimm, V., Li, Y., Rall, B.C., Rosenbaum, B. & Brose, U. (2018) Bridging Scales: Allometric Random Walks Link Movement and Biodiversity Research. Trends in Ecology & Evolution 33, 701–712.

65. Van Den Hoogen, J., Geisen, S., Routh, D., Ferris, H., Traunspurger, W., Wardle, D.A., De Goede, R.G.M., Adams, B.J., Ahmad, W., Andriuzzi, W.S., Bardgett, R.D., Bonkowski, M., Campos-Herrera, R., Cares, J.E., Caruso, T., et al. (2019) Soil nematode abundance and functional group composition at a global scale. Nature.

66. Hopkin, S.P. (1997) Biology of springtails: (Insecta: Collembola). Oxford Science Publications, Oxford.

67. Hunt, H.W., Coleman, D.C., Ingham, E.R., Ingham, R.E., Elliott, E.T., Moore, J.C., Rose, S.L., Reid, C.P.P. & Morley, C.R. (1987) The detrital food web in a shortgrass prairie. Biology and Fertility of Soils 3, 57–68.

68. Hutchinson, G.E. & Macarthur, R.H. (1959) A Theoretical Ecological Model of Size Distributions Among Species of Animals. The American Naturalist 93, 117–125.

69. Hyodo, F., Kohzu, A. & Tayasu, I. (2010) Linking aboveground and belowground food webs through carbon and nitrogen stable isotope analyses. Ecological Research 25, 745– 756.

70. Hyodo, F., Matsumoto, T., Takematsu, Y. & Itioka, T. (2015) Dependence of diverse consumers on detritus in a tropical rain forest food web as revealed by radiocarbon analysis. Functional Ecology 29, 423–429.

71. Jochum, M., Barnes, A., Brose, U., Gauzens, B., Sünnemann, M., Amyntas, A. & Eisenhauer, N. (2021) For flux’s sake: General considerations for energy-flux calculations in ecological communities. preprint, Preprints.

72. Jochum, M., Barnes, A.D., Ott, D., Lang, B., Klarner, B., Farajallah, A., Scheu, S. & Brose, U. (2017) Decreasing stoichiometric resource quality drives compensatory feeding across trophic levels in tropical litter invertebrate communities. The American Naturalist 190, 131–143.

73. Jones, C.G., Lawton, J.H. & Shachak, M. (1994) Organisms as ecosystem engineers. Oikos 69, 373–386.

74. Klarner, B., Winkelmann, H., Krashevska, V., Maraun, M., Widyastuti, R. & Scheu, S. (2017) Trophic niches, diversity and community composition of invertebrate top predators (Chilopoda) as affected by conversion of tropical lowland rainforest in Sumatra (Indonesia). PloS one 12, e0180915.

75. Krashevska, V. (2019) Changes in Nematode Communities and Functional Diversity With the Conversion of Rainforest Into Rubber and Oil Palm Plantations. Frontiers in Ecology and Evolution 7, 10.

76. Krashevska, V., Malysheva, E., Klarner, B., Mazei, Y., Maraun, M., Widyastuti, R. & Scheu, S. (2018) Micro-decomposer communities and decomposition processes in tropical lowlands as affected by land use and litter type. Oecologia 187, 255–266.

77. Krause, A., Sandmann, D., Bluhm, S.L., Ermilov, S., Widyastuti, R., Haneda, N.F., Scheu, S. & Maraun, M. (2019) Shift in trophic niches of soil microarthropods with conversion of tropical rainforest into plantations as indicated by stable isotopes (15N, 13C). PloS One 14, e0224520.

78. Laigle, I., Aubin, I., Digel, C., Brose, U., Boulangeat, I. & Gravel, D. (2018) Species traits as drivers of food web structure. Oikos 127, 316–326.

79. Lang, B., Ehnes, R.B., Brose, U. & Rall, B.C. (2017) Temperature and consumer type dependencies of energy flows in natural communities. Oikos 126, 1717–1725.

80. Larabee, F.J. & Suarez, A.V. (2015) Mandible-Powered Escape Jumps in Trap-Jaw Ants Increase Survival Rates during Predator-Prey Encounters. PLOS ONE 10, e0124871.

81. Larsen, T., Pollierer, M.M., Holmstrup, M., D’annibale, A., Maraldo, K., Andersen, N. & Eriksen, J. (2016) Substantial nutritional contribution of bacterial amino acids to earthworms and enchytraeids: A case study from organic grasslands. Soil Biology and Biochemistry 99, 21–27.

82. Lavelle, P. (1996) Diversity of soil fauna and ecosystem function. Biology International 33, 3–16.

83. Lavelle, P., Bignell, D., Lepage, M., Wolters, V., Roger, P., Ineson, P., Heal, O. & Dhillion, S. (1997) Soil function in a changing world: the role of invertebrate ecosystem engineers. European Journal of Soil Biology (France).

84. Lavelle, P. & Martin, A. (1992) Small-scale and large-scale effects of endogeic earthworms on soil organic matter dynamics in soils of the humid tropics. Soil Biology and Biochemistry 24, 1491–1498.

85. Li, Z., Tian, D., Wang, B., Wang, J., Wang, S., Chen, H.Y.H., Xu, X., Wang, C., He, N. & Niu, S. (2019) Microbes drive global soil nitrogen mineralization and availability. Global Change Biology 25, 1078–1088.

86. Luczkovich, J.J., Borgatti, S.P., Johnson, J.C. & Everett, M.G. (2003) Defining and Measuring Trophic Role Similarity in Food Webs Using Regular Equivalence. Journal of Theoretical Biology 220, 303–321.

87. Manning, P., Van Der Plas, F., Soliveres, S., Allan, E., Maestre, F.T., Mace, G., Whittingham, M.J. & Fischer, M. (2018) Redefining ecosystem multifunctionality. Nature Ecology & Evolution 2, 427–436.

88. Maraun, M., Erdmann, G., Fischer, B.M., Pollierer, M.M., Norton, R.A., Schneider, K. & Scheu, S. (2011) Stable isotopes revisited: Their use and limits for oribatid mite trophic ecology. Soil Biology and Biochemistry 43, 877–882.

89. Moore, J.C., Berlow, E.L., Coleman, D.C., Ruiter, P.C., Dong, Q., Hastings, A., Johnson, N.C., Mccann, K.S., Melville, K., Morin, P.J., & Others (2004) Detritus, trophic dynamics and biodiversity. Ecology letters 7, 584–600.

90. Moore, J.C., Mccann, K. & De Ruiter, P.C. (2005) Modeling trophic pathways, nutrient cycling, and dynamic stability in soils. Pedobiologia 49, 499–510.

91. Moore, J.C., Walter, D.E. & Hunt, H.W. (1988) Arthropod regulation of micro-and mesobiota in below-ground detrital food webs. Annual review of Entomology 33, 419–435.

92. Mougi, A. & Kondoh, M. (2016) Food-web complexity, meta-community complexity and community stability. Scientific Reports 6, 24478.

93. Mulder, C. (2006) Driving forces from soil invertebrates to ecosystem functioning: the allometric perspective. Naturwissenschaften 93, 467–479.

94. Mulder, C. & Elser, J.J. (2009) Soil acidity, ecological stoichiometry and allometric scaling in grassland food webs. Global Change Biology 15, 2730–2738.

95. Mulder, C., Den Hollander, H.A. & Hendriks, A.J. (2008) Aboveground herbivory shapes the biomass distribution and flux of soil invertebrates. PLOS ONE 3, e3573.

96. Mulder, C., Vonk, J.A., Den Hollander, H.A., Hendriks, A.J. & Breure, A.M. (2011) How allometric scaling relates to soil abiotics. Oikos 120, 529–536.

97. Newman, M.E. & Girvan, M. (2004) Finding and evaluating community structure in networks. Physical review E 69, 026113.

98. Nyffeler, M. & Birkhofer, K. (2017) An estimated 400–800 million tons of prey are annually killed by the global spider community. The Science of Nature 104.

99. Okuzaki, Y., Tayasu, I., Okuda, N. & Sota, T. (2009) Vertical heterogeneity of a forest floor invertebrate food web as indicated by stable-isotope analysis. Ecological Research 24, 1351–1359.

100. Oliverio, A.M., Geisen, S., Delgado-Baquerizo, M., Maestre, F.T., Turner, B.L. & Fierer, N. (2020) The global-scale distributions of soil protists and their contributions to belowground systems. Science Advances 6, eaax8787.

101. Ostle, N., Briones, M.J.I., Ineson, P., Cole, L., Staddon, P. & Sleep, D. (2007) Isotopic detection of recent photosynthate carbon flow into grassland rhizosphere fauna. Soil Biology and Biochemistry 39, 768–777.

102. Peschel, K., Norton, R.A., Scheu, S. & Maraun, M. (2006) Do oribatid mites live in enemy-free space? Evidence from feeding experiments with the predatory mite Pergamasus septentrionalis. Soil Biology and Biochemistry 38, 2985–2989.

103. Petchey, O.L., Beckerman, A.P., Riede, J.O. & Warren, P.H. (2008) Size, foraging, and food web structure. Proceedings of the National Academy of Sciences 105, 4191– 4196.

104. Petchey, O.L. & Belgrano, A. (2010) Body-size distributions and size-spectra: universal indicators of ecological status? Biology Letters 6, 434–437.

105. Phillips, H.R.P., Guerra, C.A., Bartz, M.L.C., Briones, M.J.I., Brown, G., Crowther, T.W., Ferlian, O., Gongalsky, K.B., Van Den Hoogen, J., Krebs, J., Orgiazzi, A., Routh, D., Schwarz, B., Bach, E.M., Bennett, J., et al. (2019) Global distribution of earthworm diversity. Science 366, 480–485.

106. Pokarzhevskii, A.D., Van Straalen, N.M., Zaboev, D.P. & Zaitsev, A.S. (2003) Microbial links and element flows in nested detrital food-webs. Pedobiologia 47, 213–224.

107. Pollierer, M.M., Dyckmans, J., Scheu, S. & Haubert, D. (2012) Carbon flux through fungi and bacteria into the forest soil animal food web as indicated by compound-specific 13C fatty acid analysis. Functional Ecology 26, 978–990.

108. Pollierer, M.M., Langel, R., Körner, C., Maraun, M. & Scheu, S. (2007) The underestimated importance of belowground carbon input for forest soil animal food webs. Ecology Letters 10, 729–736.

109. Ponge, J.-F. (2000) Vertical distribution of Collembola (Hexapoda) and their food resources in organic horizons of beech forests. Biology and Fertility of Soils 32, 508–522.

110. Potapov, A., Bellini, B., Chown, S., Deharveng, L., Janssens, F., Kováč, Ľ., Kuznetsova, N., Ponge, J.-F., Potapov, M., Querner, P., Russell, D., Sun, X., Zhang, F. & Berg, M. (2020) Towards a global synthesis of Collembola knowledge – challenges and potential solutions. Soil Organisms 92, 161–188. SOIL ORGANISMS 92(3).

111. Potapov, A., Brose, U., Scheu, S. & Tiunov, A. (2019a) Trophic position of consumers and size structure of food webs across aquatic and terrestrial ecosystems. The American Naturalist 194, 823–839.

112. Potapov, A.M., Birkhofer, K., Bluhm, S.L., Bryndova, M., Devetter, M., Goncharov, A.A., Gongalsky, K.B., Klarner, B., Korobushkin, D.I., Liebke, D., Maraun, M., Mcdonnell, R., Pollierer, M.M., Schmidt, O., Schrubovich, J., et al. ([bioRxiv]) Feeding habits and multifunctional classification of belowground consumers from protists to vertebrates. *bioRxiv*.

113. Potapov, A.M., Klarner, B., Sandmann, D., Widyastuti, R. & Scheu, S. (2019b) Linking size spectrum, energy flux and trophic multifunctionality in soil food webs of tropical land-use systems. Journal of Animal Ecology 88, 1845–1859.

114. Potapov, A.M., Korotkevich, A.Yu. & Tiunov, A.V. (2018) Non-vascular plants as a food source for litter-dwelling Collembola: Field evidence. Pedobiologia 66, 11–17.

115. Potapov, A.M., Pollierer, M.M., Salmon, S., Šustr, V. & Chen, T.-W. (2021a) Multidimensional trophic niche revealed by complementary approaches: gut content, digestive enzymes, fatty acids and stable isotopes in Collembola. Journal of Animal Ecology **n/a**.

116. Potapov, A.M., Rozanova, O.L., Semenina, E.E., Leonov, V.D., Belyakova, O.I., Bogatyreva, V.I., Degtyarev, M.I., Esaulov, A.S., Korotkevich, A.Yu., Kudrin, A.A., Malysheva, E., Mazei, Yu. A., Tsurikov, S.M., Zuev, A.G. & Tiunov, A.V. (2021b) Size compartmentalisation of energy channeling in terrestrial belowground food webs. Ecology.

117. Potapov, A.M., Scheu, S. & Tiunov, A.V. (2019c) Trophic consistency of supraspecific taxa in belowground invertebrate communities: comparison across lineages and taxonomic ranks. Functional Ecology 33, 1172–1183.

118. Potapov, A.M., Semenina, E.E., Korotkevich, A.Yu., Kuznetsova, N.A. & Tiunov, A.V. (2016) Connecting taxonomy and ecology: Trophic niches of collembolans as related to taxonomic identity and life forms. Soil Biology and Biochemistry 101, 20–31.

119. Potapov, A.M. & Tiunov, A.V. (2016) Stable isotope composition of mycophagous collembolans versus mycotrophic plants: Do soil invertebrates feed on mycorrhizal fungi? Soil Biology and Biochemistry 93, 115–118.

120. Potapov, A.M., Tiunov, A.V. & Scheu, S. (2019d) Uncovering trophic positions and food resources of soil animals using bulk natural stable isotope composition. Biological Reviews 94, 37–59.

121. Potapov, A.M., Tiunov, A.V., Scheu, S., Larsen, T. & Pollierer, M.M. (2019e) Combining bulk and amino acid stable isotope analyses to quantify trophic level and basal resources of detritivores: a case study on earthworms. Oecologia 189, 447–460.

122. Redford, K.H. (1985) Feeding and food preference in captive and wild giant anteaters (Myrmecophaga tridactyla). Journal of Zoology 205, 559–572. Wiley Online Library.

123. Rooney, N., Mccann, K., Gellner, G. & Moore, J.C. (2006) Structural asymmetry and the stability of diverse food webs. Nature 442, 265–269.

124. de Ruiter, P.C., Neutel, A.-M. & Moore, J.C. (1995) Energetics, patterns of interaction strengths, and stability in real ecosystems. Science 269, 1257.

125. de Ruiter, P.C., Van Veen, J.A., Moore, J.C., Brussaard, L. & Hunt, H.W. (1993) Calculation of nitrogen mineralization in soil food webs. Plant and Soil 157, 263–273.

126. Scheu, S. (2001) Plants and generalist predators as links between the below-ground and above-ground system. Basic and Applied Ecology 2, 3–13.

127. Scheu, S. & Setälä, H. (2002) Multitrophic interactions in decomposer food-webs. In Multitrophic level interactions pp. 223–264. Cambridge University Press, Cambridge.

128. Schmidt, O., Dyckmans, J. & Schrader, S. (2016) Photoautotrophic microorganisms as a carbon source for temperate soil invertebrates. Biology Letters 12, 20150646.

129. Schneider, K. & Maraun, M. (2009) Top-down control of soil microarthropods – Evidence from a laboratory experiment. Soil Biology and Biochemistry 41, 170–175.

130. Seppey, C.V., Singer, D., Dumack, K., Fournier, B., Belbahri, L., Mitchell, E.A. & Lara, E. (2017) Distribution patterns of soil microbial eukaryotes suggests widespread algivory by phagotrophic protists as an alternative pathway for nutrient cycling. Soil Biology and Biochemistry 112, 68–76.

131. Signorell, A. (2021) DescTools: Tools for Descriptive Statistics.

132. Soong, J.L. & Nielsen, U.N. (2016) The role of microarthropods in emerging models of soil organic matter. Soil Biology and Biochemistry 102, 37–39.

133. Stead, T.K., Schmid-Araya, J.M., Schmid, P.E. & Hildrew, A.G. (2005) The distribution of body size in a stream community: one system, many patterns. Journal of Animal Ecology 74, 475–487.

134. Steffan, S.A., Chikaraishi, Y., Dharampal, P.S., Pauli, J.N., Guédot, C. & Ohkouchi, N. (2017) Unpacking brown food-webs: Animal trophic identity reflects rampant microbivory. Ecology and Evolution 7, 3532–3541.

135. Striganova, B. (1980) Feeding of soil saprophages. Nauka, Moscow.

136. Susanti, W.I., Pollierer, M.M., Widyastuti, R., Scheu, S. & Potapov, A. (2019) Conversion of rainforest to oil palm and rubber plantations alters energy channels in soil food webs. Ecology and Evolution 9, 9027–9039.

137. Swift, M.J., Heal, O.W. & Anderson, J.M. (1979) Decomposition in terrestrial ecosystems. University of California Press, Berkeley.

138. Thakur, M.P. & Geisen, S. (2019) Trophic Regulations of the Soil Microbiome. *Trends in Microbiology*, S0966842X19301027.

139. Thakur, M.P., Phillips, H.R.P., Brose, U., De Vries, F.T., Lavelle, P., Loreau, M., Mathieu, J., Mulder, C., Van Der Putten, W.H., Rillig, M.C., Wardle, D.A., Bach, E.M., Bartz, M.L.C., Bennett, J.M., Briones, M.J.I., et al. (2020) Towards an integrative understanding of soil biodiversity. Biological Reviews 95, 350–364.

140. Thompson, R.M., Hemberg, M., Starzomski, B.M. & Shurin, J.B. (2007) Trophic levels and trophic tangles: the prevalence of omnivory in real food webs. Ecology 88, 612–617.

141. Trebilco, R., Baum, J.K., Salomon, A.K. & Dulvy, N.K. (2013) Ecosystem ecology: size-based constraints on the pyramids of life. Trends in ecology & evolution 28, 423–431.

142. Turnbull, M.S., George, P.B.L. & Lindo, Z. (2014) Weighing in: Size spectra as a standard tool in soil community analyses. Soil Biology and Biochemistry 68, 366–372.

143. Velazco, V.N., Saravia, L.A., Coviella, C.E. & Falco, L.B. (2021) Trophic resources of the edaphic mesofauna: a worldwide review of the empirical evidence. preprint, Ecology.

144. De Vries, F.T. & Caruso, T. (2016) Eating from the same plate? Revisiting the role of labile carbon inputs in the soil food web. Soil Biology and Biochemistry.

145. De Vries, F.T., Thebault, E., Liiri, M., Birkhofer, K., Tsiafouli, M.A., Bjornlund, L., Jorgensen, H.B., Brady, M.V., Christensen, S., De Ruiter, P.C., D’hertefeldt, T., Frouz, J., Hedlund, K., Hemerik, L., Hol, W.H.G., et al. (2013) Soil food web properties explain ecosystem services across European land use systems. Proceedings of the National Academy of Sciences of the United States of America 110, 14296–14301.

146. Wagg, C., Bender, S.F., Widmer, F. & Van Der Heijden, M.G.A. (2014) Soil biodiversity and soil community composition determine ecosystem multifunctionality. Proceedings of the National Academy of Sciences 111, 5266–5270.

147. Wardle, D., Bardgett, R., Klironomos, J., Setälä, H., Putten, W. & Wall, D. (2004) Ecological Linkages Between Aboveground and Belowground Biota. *Science (New York*, N.Y.) 304, 1629–1633.

148. Wardle, D.A. (2002) *Communities and ecosystems: linking the aboveground and belowground components*. Princeton University Press, Princeton and Oxford.

149. Wolkovich, E.M. (2016) Reticulated channels in soil food webs. Soil Biology and Biochemistry 102, 18–21.

150. Wolkovich, E.M., Allesina, S., Cottingham, K.L., Moore, J.C., Sandin, S.A. & De Mazancourt, C. (2014) Linking the green and brown worlds: the prevalence and effect of multichannel feeding in food webs. Ecology 95, 3376–3386.

151. Woodward, G., Ebenman, B., Emmerson, M., Montoya, J.M., Olesen, J.M., Valido, A. & Warren, P.H. (2005) Body size in ecological networks. Trends in Ecology & Evolution 20, 402–409.

152. Xu, X., Thornton, P.E. & Post, W.M. (2013) A global analysis of soil microbial biomass carbon, nitrogen and phosphorus in terrestrial ecosystems: Global soil microbial biomass C, N and P. Global Ecology and Biogeography 22, 737–749.

153. Yodzis, P. & Winemiller, K.O. (1999) In Search of Operational Trophospecies in a Tropical Aquatic Food Web. Oikos 87, 327.

